# A cadherin mutation in *Celsr3* linked to Tourette Disorder affects dendritic patterning and excitability of cholinergic interneurons

**DOI:** 10.1101/2022.03.06.483205

**Authors:** Lauren A. Poppi, K.T. Ho-Nguyen, Junbing Wu, Matthew Matrongolo, Joshua K. Thackray, Cara Nasello, Anna Shi, Matthew Ricci, Nicolas L. Carayannopoulos, Nithisha Cheedalla, Julianne McGinnis, Samantha Schaper, Cynthia Daut, Jurdiana Hernandez, Gary A. Heiman, Jay A. Tischfield, Max A. Tischfield

**Author notes:** **Corresponding author**: Max. A. Tischfield, Child Health Institute of New Jersey, Rutgers University, The State University of New Jersey, 89 French St., New Brunswick, NJ 08901.

## Abstract

*CELSR3* encodes an atypical protocadherin cell adhesion receptor that was recently identified as a high-risk gene for Tourette disorder. A putative damaging *de novo* variant was inserted into the mouse genome to generate an amino acid substitution within the fifth cadherin repeat. By contrast to *Celsr3* constitutive null animals, mice homozygous for the R774H amino acid substitution are viable and have grossly normal forebrain development. The density of cortical and striatal interneuron subpopulations is normal, but 3D geometric analysis of cortical pyramidal neurons and striatal cholinergic interneurons revealed changes to dendritic patterning and types and distributions of spines. Furthermore, patch clamp recordings in cholinergic interneurons located within the sensorimotor striatum uncovered mild intrinsic hyperexcitability. Despite these changes, *Celsr3^R774H^* homozygous mice do not show obvious ‘tic-like’ stereotypies at baseline nor motor learning impairments, but females exhibited perseverative digging behavior. Our findings show that a human mutation in *CELSR3* linked to Tourette disorder is sufficient to alter dendritic patterning in the cortex and striatum and also the intrinsic excitability of cholinergic interneurons.

## Introduction

Tourette Disorder (TD) is a childhood onset neurodevelopmental disorder associated with urges and unpleasant somatosensory phenomena, known as premonitory sensations, that serve to precipitate motor and vocal tics (Leckman et al., 2006). Although TD is traditionally classified as a movement disorder, it has common neuropsychiatric comorbidities including attention-deficit hyperactivity disorder (ADHD), obsessive compulsive disorder (OCD), and autism spectrum disorder (ASD), in addition to mood, anxiety, and sleep disorders (Hartmann and Worbe, 2018; Hirschtritt et al., 2015; Robertson, 2015; Willsey et al., 2018). TD is predicted to arise from structural and functional changes within cortico-striato-thalamo-cortical (CSTC) and basal ganglia networks that govern the planning, control, and execution of volitional motor behaviors (Draper and Jackson, 2015; Jackson et al., 2015; Kuo and Liu, 2019; Wang et al., 2011; Worbe et al., 2012).

Magnetic resonance imaging studies have suggested an association between reduced caudate volume and TD (Gerard and Peterson, 2003; Peterson et al., 2003), while post-mortem findings from a small group of adults with severe, refractory TD showed a reduction in the numbers of parvalbumin and cholinergic interneurons in the dorsal striatum (Kalanithi et al., 2005; Kataoka et al., 2010). These findings have been widely referenced to explain the neuropathogenesis of TD (Rapanelli et al., 2017a), but efforts to model these findings have produced mixed results. Focal disinhibition of the dorsal sensorimotor striatum via application of GABA_A_ receptor antagonists in rodents and non-human primates can trigger tic-like motor stereotypies (Bronfeld et al., 2013; McCairn et al., 2009; Worbe et al., 2009), whereas disinhibition in the ventral striatum leads to vocal tics (McCairn et al., 2016). Furthermore, targeted ablation of cholinergic interneurons in the rodent dorsal striatum can cause tic-like motor stereotypies following acute stress or amphetamine challenge (Xu et al., 2015), in addition to ritualistic and perseverative behaviors (Martos et al., 2017). While these animal models have provided valuable clues for understanding the pathogenesis of TD, they have notable limitations as the experimental manipulations were performed in adults, and thus they likely fail to model the types of neurodevelopmental changes found in humans.

Preclinical animal models for TD are lacking, and no true genetic models have been developed apart from *Hdc* knockout mice, whose mutation does not correspond to the inactivating point mutation found in humans (Castellan Baldan et al., 2014). This is because despite the prevalence of TD (~0.5-1% of the population) (Scharf et al., 2015), gene coding variants have been identified in only a few families, and genome-wide association studies have yielded few clues. Recently, recurrent *de novo* coding variants in several genes, including *CELSR3*, *OPA1*, and *WWC1* have been identified using whole-exome sequencing (Willsey et al., 2017). Of the identified candidates, *CELSR3* shows the strongest linkage to TD as ten different missense and/or inactivating point-mutations have been discovered to date in simplex trios and multiplex families, accounting for ~1% of clinical samples (Wang et al., 2018)(unpublished data).

*CELSR3* encodes a protocadherin cell adhesion G protein-coupled receptor that is critical for axon guidance and the development of white matter tracts that comprise CSTC circuitry (Tissir et al., 2005), and also the tangential migration of interneurons into the cortex and striatum (Ying et al., 2009). In adult animals, *Celsr3* expression is maintained in subpopulations of cortical and striatal interneurons, and also cerebellar Purkinje neurons (Ying et al., 2009). Thus, *Celsr3* expression patterns in the brain and its necessity for the development of CSTC and basal ganglia circuitry make it an attractive candidate to model TD in mice.

We have developed a genetic model for TD that expresses an analogous human amino acid substitution, R774H, within the fifth protocadherin repeat of the extracellular domain of Celsr3. To our knowledge, this represents the first genetic model for TD engineered to express the identical human mutation. Here, we investigate the impact of the Celsr3^R774H^ amino acid substitution on brain development and mouse behavior. We hypothesized that Celsr3^R774H^ mice would show perturbations to axon guidance, interneuron migration, and/or dendrite patterning. By contrast to human findings, we do not see evidence of cortical or striatal interneuron loss, and the development of major white matter tracts in the forebrain appears grossly normal. Rather, we find subtle perturbations to the structural and physiological properties of cortical and striatal neurons, including effects on dendritic patterning and membrane excitability. Using 3D pose analysis and Motion Sequencing, we do not detect overt tic-like stereotypies at baseline. Females homozygous for the R774H amino acid substitution, however, do show signs of preservative digging behavior. Our findings demonstrate that human mutations in *CELSR3* are sufficient to cause subtle but discernible changes to neuronal development and suggest the ability of neurons to functionally integrate into CSTC loops may be impaired in TD.

## Materials and methods

### Mouse lines

All experimental procedures were conducted in accordance with Rutgers Institutional Animal Care and Use Committee (IACUC) guidelines. Mice were group-housed in individually ventilated cages under a standard 12 h light/dark schedule, with controlled temperature and humidity, and *ad libitum* access to water and standard chow. Mouse lines used in this study are shown in Table S1. CRISPR/Cas9 was used within the Rutgers Gene Editing Shared Resource to produce an R774H amino acid substitution, which maps onto the fifth cadherin repeat (Fig. 1a) and corresponds to R783 in the human protein. The following single-stranded oligodeoxynucleotide template was used for targeted insertion via homology directed repair: [CAATCGGCCTGAGTTCACCATGAAAGAGTACCACCTTCGGCTCAATGAGGACGCAGCTGTAGGCACCAGTGTGGTCAGTGTGACTGCGGTAGATCACGATGCTAACAGCGCTATCAGCTACCAAATCACGGGTGGCAACACTCGGAACCGATTTGCCATC]. The following guide RNA was co-injected: [GGTAGTCGATGGTTTAGTGCCCA]. The targeted insertion added a restriction fragment length polymorphism that ablated a site recognized by Taq1 and 15 base pairs downstream of the targeted insertion. Chimeric mice were crossed with wild-type C57BL/6 animals and resulting heterozygous R774H mutant mice were backcrossed again with wild type C57BL/6 mice (Table S1) for at least three generations. The following Cre recombinase *(Drd1-Cre, A2a-Cre, Sst-Cre, Pvalb-Cre)* and reporter lines *(Celsr3-eGFP, Ai14, Chat-eGFP*, and *Pvalb*-tdT) were crossed with the *Celsr3^R774H^* line to generate double and triple transgenic lines. The *Celsr3-eGFP* knock-in mouse line was generously provided by Prof. Mario Capecchi, University of Utah, and Prof. Qiang Wu, Shanghai Jiao Tong University (Ying et al., 2009). Unless otherwise stated, all mice used in this study were young adults (P30-60).

**Figure 1.**
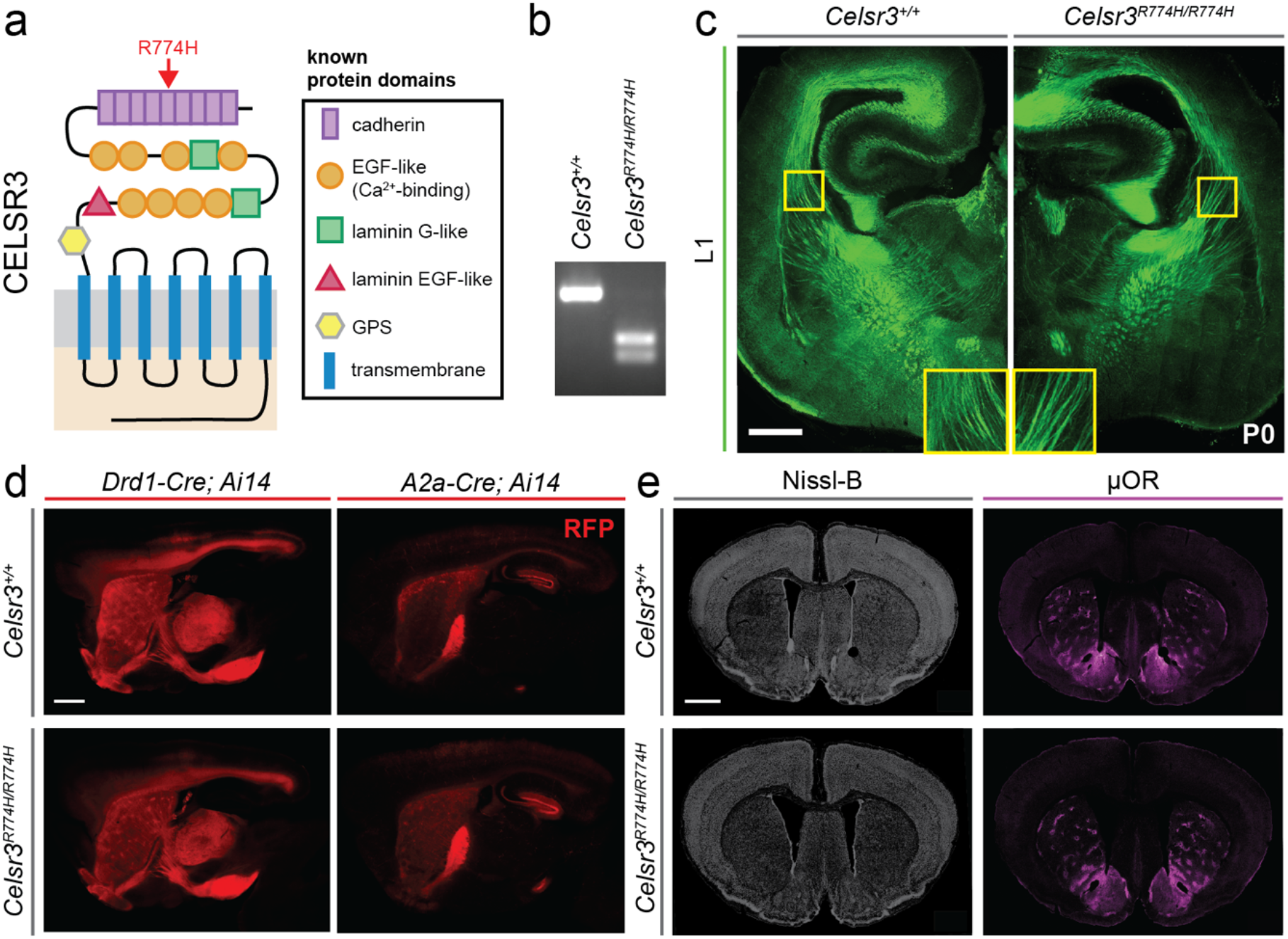
Homozygous point mutation in Celsr3 does not perturb gross organization of the mouse brain. Known domains of CELSR3 protein and location of arginine to histidine substitution (R774H, red arrow) within the fifth cadherin repeat (a). Genotyping bands for *Celsr3*^+/+^ (left) and *Celsr3^R774H/R774H^* (right; b). L1 antibody labelling in coronal sections of the P0 mouse brain shows fiber tracts in *Celsr3*^+/+^ (left) and *Celsr3^R774H/R774H^* (right) mice (c). Scale bar represents 500 μm. Sagittal view of direct pathway axon tracts in adult *Celsr3*^+/+^ (top left) and *Celsr3^R774H/R774H^* (bottom left) based on *Ai14* expression under control of *Drd1-Cre*. Sagittal view of indirect pathway fiber tracts in *Celsr3*^+/+^ (top right) and *Celsr3^R774H/R774H^* (bottom right) mice based on *Ai14* expression under control of *A2a-Cre* (d). Scale bar represents 1 mm. Nissl-B labelling shows overall cell distribution in the coronal plane at a mid-striatal position in *Celsr3*^+/+^ (top left) and *Celsr3^R774H/R774H^* (bottom left) mice. μ-OR antibody labelling shows striosomal patterning in coronal sections of *Celsr3*^+/+^ (top right) and *Celsr3^R774H/R774H^* (bottom right) striatum (e). Scale bar represents 1 mm.

### Labelling of major axon tracts

P0 mouse pups *(Celsr3^+/+^* and *Celsr3^R774H/R774H^* littermates, both sexes) were sacrificed and brains were rapidly removed and drop fixed in 4% paraformaldehyde (PFA) in 0.1 M phosphate-buffered saline (PBS) for 48 hours at 4°C. Brains were then embedded in 3% agarose and sectioned on a Leica VT1200S vibratome at 110 μm. Matched sections were labelled in parallel using rat anti-L1 (1:500, Millipore) followed by goat anti-rat Alexa Fluor 546 (1:000, Thermo Fisher). Image data were acquired on a Zeiss LSM700 confocal microscope using a 10X objective and a z-stack + tile approach. Side-by-side qualitative comparisons were made at several axial positions. Images were optimized for presentation using linear adjustments in Fiji (ImageJ).

### Labelling of direct and indirect pathways

Mice *(Drd1-Cre/+; Celsr3^+/+^;* Ai14/+, *Drd1-Cre/+; Celsr3^R774H/R774H^; Ai14/+, A2a-Cre/+; Celsr3^+/+^; Ai14/+*, and *A2a-Cre/+; Celsr3^R774H/R774H^; Ai14/+*, both sexes) were deeply anaesthetized via intraperitoneal injection of ketamine and xylazine prior to transcardial perfusion with 0.1 M PBS followed by 4% PFA in 0.1 M PBS. Brains were post-fixed in 4% PFA overnight at 4°C prior to embedding in 3% agarose and sectioning on Leica VT1200S vibratome at 120 μm. Image data were acquired on a Leica M165FC stereomicroscope with CoolLED illumination. Images were optimized for presentation using linear adjustments in Fiji (ImageJ).

### Nissl-B and mu-opiod receptor labelling

Mice *(Celsr3^+/+^* and *Celsr3^R774H/R774H^*, both sexes) were deeply anaesthetized via intraperitoneal injection of ketamine and xylazine prior to transcardial perfusion with 0.1 M PBS followed by 4% PFA in 0.1 M PBS. Brains were post-fixed overnight at 4°C prior to incubation in 30% sucrose/0.1 M PBS solution for cryoprotection and sectioning on a Leica CM1950 cryostat at 40 μm. For Nissl-B staining, matched sections were labelled in parallel with NeuroTrace 435/455 Nissl (1:500, Invitrogen). For mu-opioid receptor labelling, matched sections were labelled in parallel with rabbit anti-μOR (1:1000, immunoStar) followed by donkey anti-rabbit Alexa Fluor 647 (1:500, Thermo Fisher). Image data were acquired using a 20X objective and z-stack tile approach on a Zeiss LSM700 confocal microscope, and qualitatively compared. Images were optimized for presentation using linear adjustments in Fiji (ImageJ).

### Cortical layer markers

Mice *(Celsr3^+/+^* and *Celsr3^R774H/R774H^*, both sexes) were deeply anaesthetized via intraperitoneal injection of ketamine and xylazine prior to transcardial perfusion with 0.1 M PBS followed by 4% PFA in 0.1 M PBS. Brains were post-fixed in 4% PFA overnight at 4°C prior to incubation in 30% sucrose/0.1 M PBS solution for cryoprotection and sectioning on a Leica CM1950 cryostat at 60 μm. Matched sections were labelled in parallel with mouse anti-Satb2 (1:50, Abcam), rat anti-Ctip2 (1:1000, Abcam), and rabbit anti-Foxp2 (1:000, Abcam), followed by goat anti-mouse Alexa Fluor 546, goat anti-rat Alexa Fluor 488, and goat anti-rabbit Alexa Fluor 647 (all 1:1000, Thermo Fisher). Image data were acquired using a 20X objective and z-stack tile approach on a Zeiss LSM800 confocal microscope. Images were analysed offline in Imaris (Bitplane). Total cortical depth was measured in S1 cortex from the pial surface to the outer edge of the external capsule. Cortical layer thicknesses were measured along the same axis, guided by the fluorescent layer markers. Cortical layer thicknesses were calculated as a % of total cortical thickness. The *Spots* function was used within ROIs to determine the density and nearest neighbor distribution of labelled populations within each defined cortical layer. *Spots* data were exported into Excel (Microsoft) for further analysis. Graphing and statistical testing were done in Graphpad Prism 9. Images were optimized for presentation using linear adjustments in Fiji (ImageJ).

### Interneuron counting

Mice (*Celsr3*^+/+^, *Celsr3^R774H/R774H^*, *Sst-Cre*/+:*Celsr3*^+/+^:*Ai14*/+, *Sst*-*Cre*/+:*Celsr3^R774H/R774H^:Ai14*/+, *Celsr3*^+/+^:*Chat-eGFP* and *Celsr3^R774H/R774H^:Chat-eGFP*, both sexes) were deeply anaesthetized via intraperitoneal injection of ketamine and xylazine prior to transcardial perfusion with warm 0.1 M PBS followed by 4% PFA in 0.1 M PBS. Brains were post-fixed overnight at 4°C prior to embedding in 3% agarose and sectioning on a Leica VT1200S vibratome at either 60 μm (for parvalbumin and somatostatin interneuron counts) or 120 μm (for cholinergic interneuron counts). For parvalbumin interneuron counts (in *Celsr3^+/+^* and *Celsr3^R774H/R774H^* mice), matched sections were labelled in parallel using goat anti-parvalbumin (PV) (1:1000, Swant) followed by donkey anti-goat Alexa Fluor 488 (1:1000, Thermo Fisher). For somatostatin interneuron counts (In *Sst-Cre/+; Celsr3^+/+^; Ai14/+* and *Sst-Cre*/+; *Celsr3^R774H/R774H^*; *Ai14*/+ mice), matched sections were labelled in parallel using rabbit anti-RFP (1:1000, Rockland) followed by donkey anti-rabbit Alexa Fluor 546 (1:1000, Thermo Fisher). For cholinergic interneuron counts, matched sections were labelled in parallel using chicken anti-GFP (1:500, Aves Labs) and goat anti-choline acetyltransferase (ChAT) (1:200, Millipore) followed by donkey anti-chicken Alexa Fluor 488 and donkey anti-goat Alexa Fluor 546 (both 1:1000, Thermo Fisher). Image data were acquired using a 20X objective and z-stack tile approach with a maximum step size of 2 μm on a Zeiss LSM 700 confocal microscope. Images were analysed offline and blinded to genotype in Fiji (Image J). Interneuron counts were quantitatively compared at 4 predefined anterio-posterior axis positions relative to bregma: position 1 (1.53 to 0.85 mm), position 2 (0.85 to 0.13 mm), position 3 (0.13 to −0.59 mm), and position 4 (−0.59 to −1.31 mm) (Franklin and Paxinos, 2012). Graphing and statistical testing were done in Graphpad Prism 9.

### Viral sparse cell labeling

Mice were anesthetized with 1-3 % vaporized isoflurane in oxygen (1 L/min) and placed on a stereotaxic frame. *Pvalb-Cre/+; Celsr3^R774H/R774H^* animals and *Pvalb-Cre/+; Celsr3^+/+^* littermate control animals were injected with a cocktail of 2 adenoviruses (AAV9-TRE-DIO-vCre and AAV9-TRE-vDIO-GFP-tTA) diluted in sterile saline (1:1:18 ratio of AAV9-TRE-DIO-vCre to AAV9-TRE-vDIO-GFP-tTA to 0.9% NaCl) bilaterally into S1 (+/− 1.80 ML, 0.00 AP, −1.75DV; 500 nL each injection at a rate of 100 nL/min). This sparse labelling system, provided by Dr. Minmin Luo, Tsinghua University, consists of a controller vector that contains a Tetracycline Response Element promoter (TRE) and a Cre-dependent expression cassette (double-floxed inverse open reading frame) encoding a mutated Cre-recombinase (vCre) that only recognizes vLoxP sites (Lin et al., 2018). The amplifier vector contains a vCre-dependent expression cassette encoding membrane-anchored GFP (mGFP) and the tetracycline-controlled transactivator (tTA) downstream of an internal ribosome entry site. When these viruses are injected into mice that express Cre-recombinase, vCre is flipped into the correct reading frame. vCre can then flip the amplifier expression cassette into the correct orientation, resulting in GFP and tTA expression. Under basal conditions, the TRE promoter is “leaky” and provides very low levels of vCre expression, and only a few neurons will produce enough vCre to flip the amplifier expression cassette into the right orientation. In these sparsely populated neurons, tTA can bind to the TRE promotor on both the control and amplifier vectors, boosting mGFP expression in a positive feedback loop. Following surgery, buprenorphine SR (1.5 mg/kg), carprofen (5 mg/kg) and sterile saline were administered for 3 days post-surgery and the health and welfare of mice were closely monitored. 3 weeks post-surgery, mice were transcardially perfused as described above.

### Anatomical recovery of cortical pyramidal neurons

Fixed brains were embedded in 3% agarose blocks and sectioned on a Leica VT1000 vibratome at 110 μm. mGFP signal was amplified by incubating in chicken anti-GFP (1:500, Aves Labs) followed by goat anti-chicken Alexa Fluor 488 (1:1000, Thermo Fisher). mGFP expressing cells were imaged on a Leica LSM700 confocal microscope at 20X using a z-stack tile approach with maximum z-steps of 1 μm. For spine counts, secondary dendrites were imaged on a Leica LSM800 confocal microscope using a 63X oil immersion lens with minimum z-step distance (0.46 μm) and post-hoc deconvolution. z-stack tile images were imported into Imaris (RRID:SCR_007370) and neurites were semi-automatically traced using the autodepth feature in *Filaments*. Tracing was performed independently by two different experimenters and blinded to mouse genotype. Soma were rendered using *Surfaces* for illustration purposes only. Spines were detected semiautomatically, and diameters were recomputed using the shortest distance from distance map algorithm. Spines were classified into 4 distinguished classes: stubby, mushroom, long thin, and filopodia using the *ClassifySpines* Xtension and specified criteria (Table S2). All *Filaments* and *ClassifySpines* data were exported into Excel for further analysis, and graphic and statistical testing were done in GraphPad Prism 9 (RRID:SCR_002798).

### Electrophysiology

Mice *(Celsr3^+/+^; Chat-eGFP* and *Celsr3^R774H/R774H^; Chat-eGFP*, both sexes) were anesthetized with an intraperitoneal injection of ketamine + xylazine prior to rapid decapitation and brain dissection (de Oliveira et al., 2010). Coronal 300 μm sections were taken on a Leica VT1200S vibratome in ice-cold sucrose substituted cerebrospinal fluid (aCSF) containing (in mM): 250 sucrose, 25 NaHCO_3_, 10 glucose, 2.5 KCl, 1 NaH_2_PO_4_, 1 MgCl and 2.5 CaCl_2_. Ringers’ solutions were continually bubbled with 95% O_2_ / 5% CO_2_ to maintain oxygenation and neutral pH. Sections were allowed to recover for 1 hour at room temperature in normal aCSF (118 mM NaCl substituted for sucrose) prior to recording. aCSF was continually bubbled with 95% O_2_ / 5% CO_2_. Evoked action potential characterization was done using a potassium gluconate based internal solution containing (in mM): 135 K.gluconate, 8 NaCl, 10 HEPES, 0.1 EGTA, 0.3 Na_3_GTP, and 2 Mg_2_ATP. Biotin hydrobromide (0.2%, Biotium) was added to the internal solution. Data were amplified using a Multiclamp 200B amplifier, digitized using a Digidata 1550A, and acquired using pClamp11 (Molecular Devices, RRID:SCR_011323). Series resistance (R_s_), membrane resistance (R_m_), membrane capacitance (C_m_), and resting membrane potential (RMP) were measured at the beginning of recording and monitored throughout. Evoked AP characteristics were recorded within 1 min of membrane breakthrough. Bridge balances were applied in current clamp mode. Voltages have not been corrected for liquid junction potential. Cholinergic interneurons in the dorsolateral striatum were fluorescence targeted via their expression of eGFP, and their identities were confirmed physiologically via relatively depolarized RMPs (~ −55mV), prominent voltage sag, slow AHP currents, and relatively wide action potential waveforms. AP threshold was measured using the first derivative of the AP and was defined as the voltage at which dV/dt = 10 mV.s^-1^. Data were excluded from analysis if R_s_ > 30 MOhm or if △Rs > 20% over the course of the recording. Electrophysiology data were analyzed offline in Axograph X (Axograph, Sydney, RRID:SCR_014284). At the end of recording, slices were dropped into 4% PFA for post-hoc anatomical recovery. Slices were kept in 4% PFA overnight at 4°C, washed in 0.1 M PBS, then stored in 0.1 M PBS + 5 mM NaN_3_ at 4°C until further processing.

### Anatomical recovery of striatal cholinergic interneurons

Slices were incubated in in Alexa-546 conjugated streptavidin (1:50, Thermo Fisher) for 2 hours at room temperature. Slices were then mounted onto slides in Fluoromount-G (Southern Biotech). Biotin-filled and fluorescently tagged cholinergic interneurons were imaged on a Zeiss LSM700 confocal microscope using a 63X Apo-plan oil immersion lens and a z-stack + tile approach. Labelled cell somata were checked for eGFP expression to confirm cholinergic identity. Step size was set to minimum (0.46 μm). Raw .czi data were converted and imported into Imaris (Bitplane) and neurites were traced using *Filaments* as described above. Soma volumes were rendered for presentation purposes using *Surfaces*, but due to potential perturbation from recording, soma sizes were not compared. Spines were also measured on ROI images using *ClassifySpines*, as described above. Fractal dimension (D_B_) and lacunarity of cholinergic interneuron neurites were measured using the FracLac ImageJ plugin (http://rsb.info.nih.gov/ij/plugins/fraclac/FLHelp/Introduction.htm). Data were exported from Imaris and ImageJ to excel for further analysis, and graphing and statistical testing were done in Graphpad Prism9.

### Marble burying assay

Mice *(Celsr3^+/+^* and *Celsr3^R774H/R774H^*, both sexes) were gently placed into a rectangular arena with a 5 cm base of Beta Chip bedding (Northeastern Products), where 20 glass marbles had been placed on top of the bedding in a 4 x 5 grid pattern. After 30 minutes, the mouse was returned to their home cage and the number of marbles buried were counted. A marble was counted as ‘buried’ if it was buried 50% or more. This assay was repeated over three consecutive days and the number of marbles buried across the three trials was averaged for each animal.

### 3D depth imaging and pose analysis

Data was acquired and processed as previously described (Bohic et al., 2021). For motion sequencing image data acquisition, mice were gently placed into a 17” diameter cylindrical enclosure with 17”-high walls (US Plastics) and allowed to roam freely for 20 minutes while being recorded with a Kinect2 depth-sensing camera (Microsoft). Depth data were modelled as described previously (Wiltschko et al., 2015). Raw depth frames were collected at 30 Hz using custom C# code. Frames were 512 x 424 pixels and each pixel carried a 16-bit integer value denoting the distance from the sensor in mm. Frames were compressed and analyzed offline. Briefly, the mouse’s center and orientation were determined using an ellipse fit. Then, an 80 x 80 pixel box was drawn around the mouse, and the mouse was rotated to face the right hand side. Next, if the tracking model was used, missing pixels were identified by their likelihood according to the Gaussian model. Low-likelihood pixels were treated as missing data and principal components (PCs) are computed using probabilistic PCA (Roweis, 1998; Tipping and Bishop, 1999). Finally, frames were projected onto the first 10 PCs, forming a 10-dimensional time series that described the mouse’s 3D pose trajectory. These data were used to train an autoregressive hidden Markov model (AR HMM) with 3 lags to cluster mouse behavioral dynamics into discrete ‘modules’ with state number automatically identified using a hierarchical Dirichlet process. Each state was comprised of a vector autoregressive process that captures the evolution of the 10 PCs over time. The model was fit using Gibbs sampling as described in Wiltschko et al. (2015) using freely available software (https://github.com/mattjj/pybasicbayes). Model output was insensitive to all but two hyperparameters, which were set using unsupervised techniques for determining the length scales for discrete behaviors as was previously published (Wiltschko et al., 2015).

## Results

### Gross phenotypic presentation of Celsr3^R774H^ mice

*Celsr3*^*R774H*/+^ and *Celsr3^R774H/R774H^* animals on a pure C57BL/6 background were born at normal Mendelian ratios, had normal weights, and were indistinguishable by eye from littermate controls. By contrast, *Celsr3* constitutive null animals are perinatal lethal, suggesting the Celsr3^R774H^ mutant protein still retains key functions. No obvious tics or motor stereotypies were apparent at baseline by eye, and there were no signs of hair loss or skin lesions indicative of compulsive grooming behavior. We focused our remaining investigation on animals that were homozygous for the amino acid substitution.

### Axon tract development is grossly normal in Celsr3^R774H^ mice

Celsr3 is required for the development and guidance of major forebrain axon tracts, such as the anterior commissure and internal capsule, which contains corticostriatal, thalamocortical, and corticothalamic axons. Gross anatomy and overall size of *Celsr3^R774H/R774H^* mouse brains appeared normal. Antibody labelling against neuronal cell adhesion protein L1 in embryonic day (E)18.5 brain sections showed that the development and trajectories of major forebrain axon tracts in the internal capsule, anterior commissure, and corpus callosum were normal in *Celsr3^R774H/R774H^* animals (Fig. 1C). Celsr3 is also required for the development of key basal ganglia pathways, including axonal projections from areas such as the striatum, subthalamic nucleus, and the substantia nigra pars compacta to the globus pallidus (Jia et al., 2014). Striatonigral axons in the direct pathway terminating in the globus pallidus internus (GPi) and substantia nigra (SNr) were visualized by crossing *Drd1a-Cre* and *R26:Ai14* reporter lines (Table S1). Striatonigral fiber tracts showed normal development in adult *Drd1a-Cre;Celsr3^R774H/R774H^;R26:Ai14* mice and terminated appropriately in the GPi (n = 3, Fig. 1D). Next, we crossed *A2a-Cre* and *R26:Ai14* reporter lines (Table S1) to visualize striatopallidal axons that terminate in the globus pallidus externus (GPe). We did not detect any qualitative differences in the pattern of tdTomato-positive fibers terminating in the GPe of *Drd1a-Cre;Celsr3^R774H/R774H^;R26:Ai14* animals (n = 4) compared to littermate controls (n = 3, Fig. 1D). We also did not observe instances of wandering axons or misinnervation by direct and indirect pathway axons. Overall cellular organization within the cortex and subcortical structures was comparable between *Celsr3*^+/+^ (n = 2) and *Celsr3^R774H/R774H^* mice (n = 2) based on Nissl-B staining (Fig. 1E). In the striatum, the formation of the matrix and striosome compartments also appeared normal in *Celsr3^R774H/R774H^* animals according to the pattern of mu-opioid receptor labelling (n = 2, Fig. 1E). Thus, this Celsr3^R774H^ amino acid substitution within the fifth cadherin repeat does not affect the ability of the protein to regulate axon guidance in the forebrain in a manner that is detectable with the qualitative anatomical techniques used.

### Celsr3^R774H^ mice have organized cortical layering and do not show interneuron loss

Cortical layering, as assessed by TBR1, CTIP2, and SATB2 immunostaining, was normal in *Celsr3^R774H/R774H^* animals (n = 3) compared to littermate controls (n = 3, Fig. 2A). The relative radial thickness of each cortical layer was also normal in *Celsr3^R774H/R774H^* animals (n = 3, Fig. 2B, p = 0.9742, Chi-square test), and nearest neighbor analysis showed normal distribution of labelled cortical neurons (Fig. 2B, p = 0.2275, 2way ANOVA). *Celsr3* is expressed by E13.5 in the ganglionic eminences, which give rise to cortical and striatal interneurons, and has been reported to regulate the tangential migration of cortical interneurons (Ying et al., 2009). Immunolabeling against parvalbumin showed the density of cortical parvalbumin interneurons was normal in *Celsr3^R774H/R774H^* mice (Fig. 2D). Using *somatostatin-Cre* and the *R26:Ai14* reporter line to lineage label somatostatin interneurons, there were also no differences in the density of these interneurons in the cortex of *Celsr3^R774H/R774H^* mice (Fig. 2F). Thus, cell proliferation and the radial and tangential migration of cortical pyramidal neurons and interneurons, respectively, was unaffected in *Celsr3^R774H/R774H^* animals.

**Figure 2.**
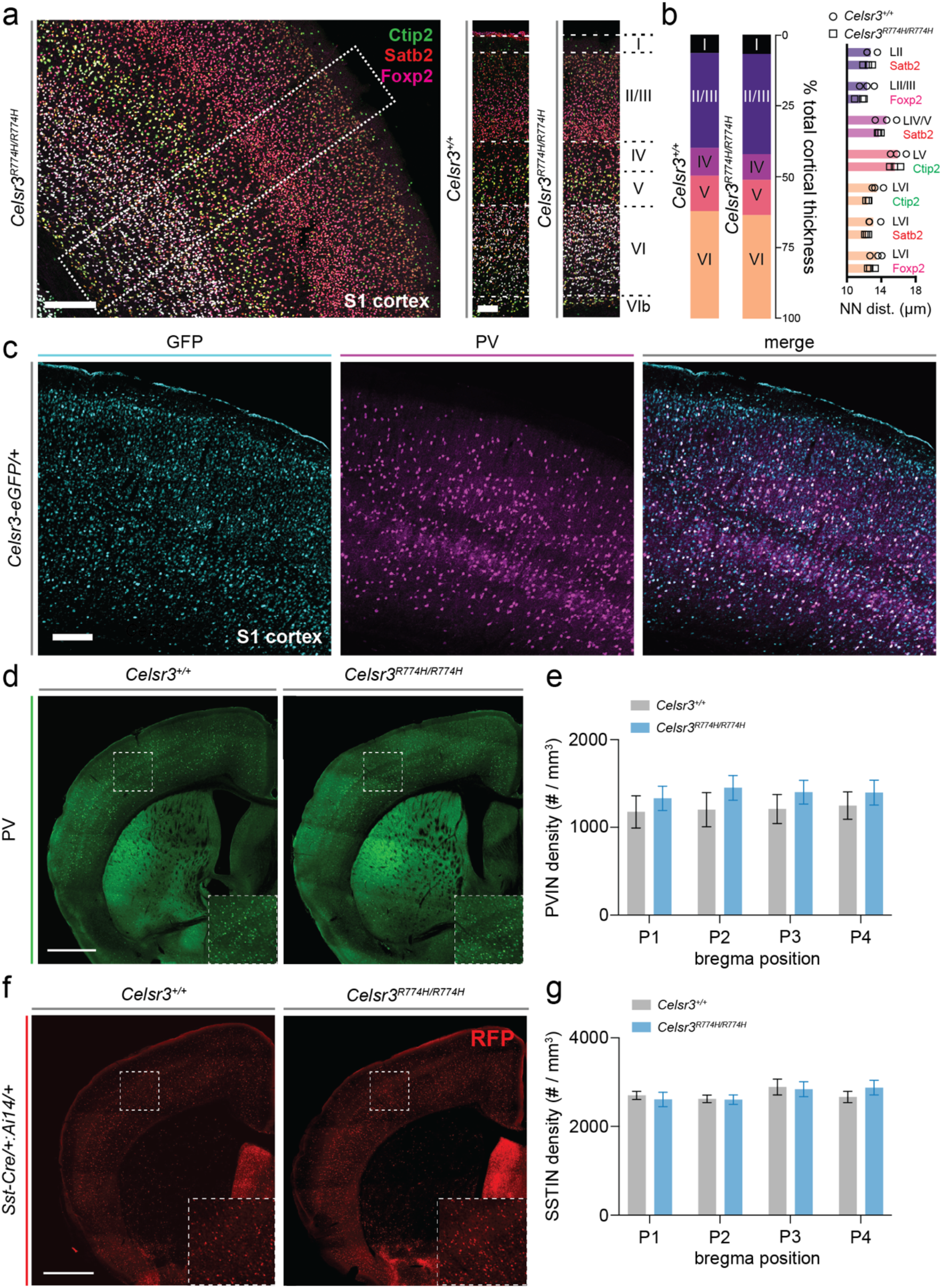
Cortical layering and inhibitory interneuron patterning is not significantly impacted by the R774H amino acid substitution in Celsr3. Representative image of cortical layers in a *Celsr3^R774H/R774H^* mouse somatosensory (S1) cortex (large image, a). Scale bar represents 200 μm. Representative ROIs of *Celsr3*^+/+^ and *Celsr3^R774H/R774H^* (smaller images), with layer positions I-VI marked (a). Scale bar represents 100 μm. Relative cortical layer thickness in *Celsr3*^+/+^ (left bar, n = 3) and *Celsr3^R774H/R774H^* (right bar, n = 3, p = 0.9742, Chi-square test, b). Nearest neighbor distances across labelled populations within defined layers (p = 0.2275, 2way ANOVA, b). Celsr3-eGFP expression in mouse S1 cortex co-labelled for parvalbumin (PV, c). Scale bar represents 200 μm. Representative images of cortical parvalbumin interneurons (PVINs) in *Celsr3*^+/+^ (left) and *Celsr3^R774H/R774H^* (right) mice (d). Scale bar represents 1 mm. Comparison of cortical PVIN density at four different AP positions *(Celsr3^+/+^* n = 8, *Celsr3^R774H/R774H^* n = 7, p = 0.4159, 2way ANOVA, e). Representative images of cortical somatostatin interneurons (SSTINs) in *Sst-Cre/+:Celsr3^+/+^:Ai14*/+ (left) and *Sst-Cre*/+:*Celsr3^R774H/R774H^:Ai14*/+ (right) mice (f). Scale bar represents 1 mm. Comparison of cortical SSTIN density at four different AP positions (p = 0.8944, 2way ANOVA, g).

### Cortical pyramidal neuron dendritic patterning is affected in Celsr3^R774H/R774H^ mice

We examined *Celsr3* expression in the cortex using a green fluorescent protein (GFP) knock-in reporter line *(Celsr3^GFP^)* that faithfully recapitulates its endogenous expression patterns (Ying et al., 2009). GFP labelling shows expression is maintained in subsets of parvalbumin interneurons in juvenile and adults (Fig. 2C). Celsr3 is required for neurite development and dendritic patterning in cortical pyramidal and hippocampal CA1 neurons (Feng et al., 2012; Zhou et al., 2010), so we examined whether the dendritic arborizations of cortical parvalbumin interneurons were properly patterned in *Celsr3^R774H/R774H^* mice using a Cre-dependent viral sparse cell labelling approach to mark parvalbumin (PV) interneurons with GFP (Lin et al., 2018). We crossed a *PV-2A-Cre* allele onto the *Celsr3^R774H/R774H^* background and injected the virus into the somatosensory cortex. Most labelled neurons were located in deep layer 5 of the cortex but surprisingly, most were not positive for parvalbumin immunostaining. Instead, these neurons had typical cortical pyramidal neuron morphology with basal and long apical dendrites. Crossing these animals to the *R26:Ai14* reporter line showed diffuse td-Tomato expression throughout the cortex, suggesting the *PV-2A-Cre* allele went germline, consistent with previous reports (Luo et al., 2020). Nonetheless, 3D neuronal reconstructions revealed that the basal dendrites of *Celsr3^R774H/R774H^* deep layer 5 pyramidal neurons were less arborized than littermate controls (Fig. 3B). Basal dendrites were also analyzed separately by excluding apical branches from the dataset (Fig. 3C). Sholl analysis revealed a genotype effect for the complexity of *Celsr3^R774H/R774H^* pyramidal neuron basal dendrites (Fig. 3D; *Celsr3*^+/+^ n = 6; *Celsr3^R774H/R774H^* n = 8; two-way ANOVA genotype effect p < 0.001). The area under the Sholl curve was 1639 +/− 40.05 and 1204 +/− 27.62 for *Celsr3*^+/+^ and *Celsr3^R774H/R774H^*, respectively. There was also a significant genotype effect when comparing branch depth, which reflects the number of times a dendrite has branched since leaving the soma, with total length (Fig. 3E; p = 0.0271, two-way ANOVA). There was no significant difference in the number of branch points *(Celsr3^+/+^* = 16.33 +/− 2.81, *Celsr3^R774H/R774H^* = 15.22 +/− 1.52, p = 0.7110, unpaired t test) or dendritic straightness *(Celsr3^+/+^=* 0.9411 +/− 0.003, *Celsr3^R774H/R774H^* = 0.9298 +/− 0.006, p = 0.1805, unpaired t test). We also did not see evidence of increased number of self-crossings. There was no difference in the density of spines along the secondary basal dendrites between *Celsr3*^+/+^ (8.60 / 10 μm) and *Celsr3^R774H/R774H^* (8.94 / 10 μm). However, when spines were classified according to morphology *(e.g*. stubby, mushroom, long-thin, filopodia), and the relative densities were compared using the *ClassifySpines* IMARIS plug-in, the proportion of stubby and long-thin spines detected along a single length of dendrite appeared shifted in *Celsr3^R774H/R774H^* mice (Fig. 3G). There was also a significant reduction in stubby spines in *Celsr3^R774H/R774H^* animals (p = 0.033), and a trend toward an increase in long thin spines (p = 0.055, t-tests with multiple comparison correction). Thus, the Celsr3^R774H^ amino acid substitution is sufficient to alter dendritic patterning and the types and distributions of spines in deep layer cortical pyramidal neurons.

**Figure 3.**
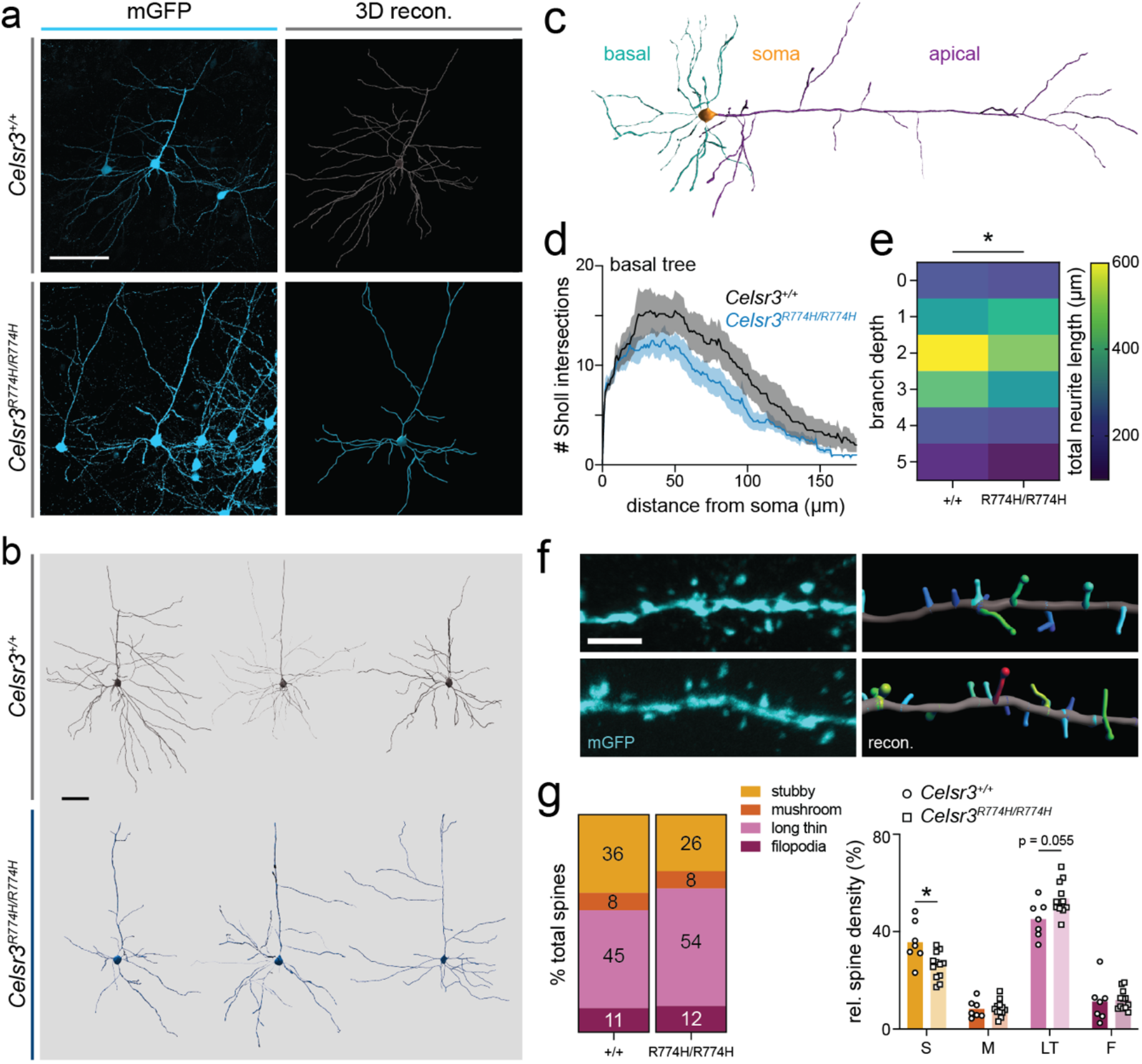
Basal dendrites of *Celsr3*-mutant cortical pyramidal neurons show reduced complexity. Representative images of confocal images (left) and their 3D reconstructions (right, a). Scale bar represents 100 μm. Representative reconstructions of *Celsr3*^+/+^ (top, grey) and *Celsr3^R774H/R774H^* (bottom, blue) cortical pyramidal neurons (b). Scale bar represents 50 μm. Schematic showing denotation of basal dendrites (blue) versus apical dendrites (purple, c). Sholl analysis of *Celsr3*^+/+^ (n = 6, black) and *Celsr3^R774H/R774H^* (n = 8, blue) basal dendrites (genotype effect p < 0.001, 2way ANOVA, d). Shaded area represents SEM. Heat map comparing total neurite length vs. branch depths (p = 0.0271, 2way ANOVA, e). Representative confocal images of secondary dendrites (left) and their 3D reconstruction and classification (right, f). Scale bar represents 2 μm. Relative spine density by class: stubby (S), mushroom (M), long thin (LT) and filopodia (F) in *Celsr3*^+/+^ and *Celsr3^R774H/R774H^* mice (stubby spines p = 0.03, long thin spines p = 0.055, multiple Holm-Šídák t-test with multiple comparison correction, g).

### Disorganization of striatal cholinergic interneuron neurite patterning

We examined neurite patterning of single striatal cholinergic interneurons using biotin filling during recording and post-hoc anatomical recovery. *Celsr3^R774H/R774H^* (n = 13) cholinergic interneurons showed increased neurite complexity compared to *Celsr3*^+/+^ cholinergic interneurons (n = 6, p < 0.001, 2way ANOVA, Fig. 5b). The area under the curve values for cholinergic interneuron Sholl plots were 2709 +/− 44.2 and 3165 +/− 41.1, for *Celsr3*^+/+^ and *Celsr3^R774H/R774H^*, respectively, and each dataset fell outside the 95% confidence interval of the opposing genotype. Data presented in heatmap form allowed for alternative visualization of relative complexity of neurites with increasing distance from the soma, and Celsr3^R774H/R774H^ neurites appear to be more compact on average (Fig. 5c). The number of branch points trended towards a significant increase in *Celsr3^R774H/R774H^* cholinergic interneurons (p = 0.06, t-test, Fig. 5d), and neurite straightness trended towards a significant decrease in *Celsr3^R774H/R774H^* cholinergic interneurons (p = 0.05, t-test, Fig. 5e). As these results may not have fully explained either the Sholl effect or the striking visual appearance of *Celsr3^R774H/R774H^* cholinergic interneurons, we analysed the fractal geometry and lacunarity of their neurite patterns in 2D. While fractal dimension (D_B_) was similar in *Celsr3*^+/+^ and *Celsr3^R774H/R774H^* cholinergic interneurons (p = 0.2636, Mann-Whitney test), lacunarity was significantly increased in *Celsr3^R774H/R774H^* compared to controls (p = 0.0379, t-test).

### Striatal cholinergic interneuron physiology

Dorsolateral striatal cholinergic interneurons of both *Celsr3*^+/+^ and *Celsr3^R774H/R774H^* mice had characteristically large somata, and upon membrane breakthrough, had a relatively depolarized resting membrane potential (RMP). Passive membrane properties were not significantly different between *Celsr3*^+/+^ and *Celsr3^R774H/R774H^* mice. Membrane impedance (Rm) was 184.8 +/− 8.82 MOhm and 205.1 +/− 11.89 MOhm for *Celsr3*^+/+^ (n = 31) and *Celsr3^R774H/R774H^* (n = 39) Cholinergic interneurons, respectively (p = 0.4238, Mann-Whitney test). Membrane capacitance (C_m_) was 33.52 +/− 1.10 pF and 34.05 +/− 0.97 pF for *Celsr3*^+/+^ and *Celrs3^R774H/R774H^* cholinergic interneurons, respectively (p = 0.7214, t-test). Membrane time constant (tau) was 2.88 +/− 0.16 ms and 2.91 +/− 0.16 ms for *Celsr3*^+/+^ and *Celsr3^R774H/R774H^* cholinergic interneurons, respectively (p = 0.8870, t-test). Resting membrane potential (RMP) was on average more depolarized in *Celsr3^R774H/R774H^* cholinergic interneurons (p = 0.037, t-test, Fig. 6b). Rheobase (minimum current injection step required to elicit an action potential) was not significantly affected (p = 0.3505, Mann-Whitney test, Fig. 6e). The action potential (AP) threshold was significantly more depolarized in *Celsr3^R774H/R774H^* cholinergic interneurons (p = 0.0456, t-test, Fig. 6f). The f/I plots for *Celsr3*^+/+^ (n = 29) and *Celsr3^R774H/R774H^* (n = 25) required different nonlinear fits (p < 0.001, Fig. 6g). This indicated a tendency for *Celsr3^R774H/R774H^* cholinergic interneurons to fire at a higher frequency in response to somatic current injection compared with *Celsr3*^+/+^ cholinergic interneurons. AP frequency was significantly higher in *Celsr3^R774H/R774H^* compared to *Celsr3*^+/+^ cholinergic interneurons with 200 pA current injection (p = 0.038, t-test, Fig. 6g). Thus, Celsr3^R774H^ is sufficient to alter the membrane properties of cholinergic interneurons.

### *Celsr3^R774H/R774H^* female mice show preservative digging behavior

The Celsr3^R774H^ variant is reported in an individual with TD and comorbid ADHD. We assessed activity levels in SmartCages in mixed sex cohorts using infrared beam breaks. *Celsr3^R774H/R774H^* mice did not show an increase in overall activity compared to littermate controls (Extended Data 1, p = 0.6249, 2way ANOVA). *Celsr3^R774H/R774H^* mice were trending towards a significant increase in the number of vertical rears (upper IR beam break) compared to controls (Extended Data 1, p = 0.0717, 2way ANOVA). *Celsr3^R774H/R774H^* mice showed normal latency (p = 0.6300, 2way ANOVA) and speed progression (p = 0.6760, 2way ANOVA) on an accelerated rotarod, suggesting motor coordination and learning is intact in these animals (Extended Data 1). We examined perseverative behaviors using the marble burying assay, *i.e*. counting the number of marbles buried within a 30 minute span (see schematic Fig. 6A). Male *Celsr3*^+/+^ mice (n = 11) buried a similar number of marbles (14.30 +/− 1.00) to *Celsr3^R774H/R774H^* mice (n = 8, 14.59 +/− 1.10, p = 0.8504, two-tailed t test). However, female *Celsr3^R774H/R774H^* mice (n = 11) buried a significantly higher number of marbles on average (16.66 +/− 0.64) compared to littermate *Celsr3*^+/+^ controls (n = 13, 13.92 +/− 0.72, p = 0.0105; two-tailed t test, Fig. 6B).

### Celsr3^R774H^-mutant mice do not show tic-like stereotypies at baseline

To determine whether *Celsr3^R774H/R774H^* mice exhibit motor stereotypies, we used Motion Sequencing to analyze motor behavior while animals explored an open field (circular diameter 17 inches, Fig. 6c) for 20 minutes (Bohic et al., 2021; Wiltschko et al., 2015). MoSeq can learn to identify stereotyped motor modules (e.g. rear, groom, scrunch) and calculate the usage frequencies, as well as transition probabilities, that determine how these modules are assembled into short motion sequences. Single module usage frequencies were not significantly different between *Celsr3*^+/+^ and *Celsr3^R774H/R477H^* mice after strict multiple correction testing (Fig. 6e). We saw numerous, albeit subtle, changes to first order transition probabilities that link motor modules together, but action selection did not appear to be appreciably affected according to the overall rate of entropy (Extended Data 3) (Markowitz et al., 2018). Next, we used computational methods based on natural language processing to examine how these short sequences were grouped into larger embeddings (Bohic et al., 2021). Interestingly, this method showed that Celsr3^R774H^ animals could be distinguished from littermate controls with an accuracy rate of 81%, which was greater than what was seen according to module usages or first order transition probabilities alone. Thus, this data suggests that Celsr3^R774H^ animals do have subtle changes to motor behavior that are reflected by how short sequences are embedded into larger action sequences over longer timescales based on higher order transition probabilities. Finally, we used depth imaging data to calculate time spent in the center versus the periphery of the circular open field. We did not detect any significant changes to the amount of time spent in the center, suggestion that overall levels of anxiety were unaffected in Celsr3^R774H^ animals (Extended Data 4).

## Discussion

We present a phenotypic analysis of a mouse model for Tourette Disorder engineered to express a putative damaging variant in *Celsr3* that causes an amino acid substitution within the fifth extracellular cadherin repeat. To our knowledge, this is the first model for Tourette Disorder that expresses the identical human mutation. Putative damaging variants in *Celsr3* identified to date include missense mutations and a frameshift that leads to a stop-gain in the second laminin-G like domain (Wang et al., 2018). The latter suggests TD associated variants in *Celsr3* exert loss-of-function effects on the protein. By contrast with *Celsr3* constitutive null animals that die at birth (Tissir et al., 2005), animals homozygous for the Celsr3^R774H^ amino acid substitution were viable and fertile. This suggests the mutation may exert only mild, partial loss-of-function effects on the protein, although gain-of-function effects may be possible as well.

Constitutive loss of *Celsr3* affects axon guidance and the development of white matter tracts in the internal capsule, including corticostriatal, corticothalamic, and thalamocortical fibers that comprise CSTC pathways (Tissir et al., 2005; Zhou et al., 2008). While Celsr3 is required cell autonomously for corticospinal and corticostriatal axon pathfinding, is thought to guide thalamocortical and corticothalamic axons in a non-cell autonomous manner via its activity in guidepost neurons (Zhou et al., 2008). In addition, Celsr3 is required for the formation of axon tracts within basal ganglia circuits (Jia et al., 2014). By contrast with *Celsr3* constitutive null animals, the Celsr3^R774H^ amino acid substitution modelled in the present study does not cause appreciable misrouting of axons (Fig. 1). More subtle changes to the abilities of axons to terminally branch and/or synapse appropriately onto neurons may be present and functionally significant and will need to be investigated further.

*Celsr3* is expressed by ~E14.5 in the mouse ganglionic eminences, which produce cortical and striatal interneurons (Tissir and Goffinet, 2006). The role of Celsr3 in the tangential migration of interneurons from the preganglionic eminences has been debated (Feng et al., 2012). Constitutive loss of protein function in homozygous *Celsr3^GFP^* knock-in mice is reported to affect tangential interneuron migration (Ying et al., 2009). Cortical interneuron loss is reported in these animals as tangentially migrating calretinin-expressing interneurons appear to become trapped at the boundary between the cortex and the striatum. An increase of calretinin expressing interneurons was reported in the striatum, and they were abnormally distributed compared to control animals (Ying et al., 2009). Reports of *Celsr3*-mediated alterations in interneuron migration are intriguing in light of findings from post-mortem brains of adults with severe TD that showed loss of striatal parvalbumin and cholinergic interneurons (Kalanithi et al., 2005; Kataoka et al., 2010). This prompted us to examine whether similar interneuron deficits may be present in the brains of homozygous Celsr3^R774H^ animals. We do not detect any significant loss of cortical parvalbumin or somatostatin interneurons (Fig. 2), nor do we see loss of striatal parvalbumin, somatostatin, or cholinergic interneurons (Fig. 4). In agreement, *Hdc* knockout mice, which have been used to model a familial stop-gain mutation identified in TD, have normal cortical layering and also do not show signs of parvalbumin or cholinergic interneuron loss in the cortex and/or striatum (Abdurakhmanova et al., 2017). Thus, our findings and those in *Hdc* knockout animals seem to suggest that striatal interneuron loss may be present only in rarer and more severe cases across the TD spectrum.

**Figure 4.**
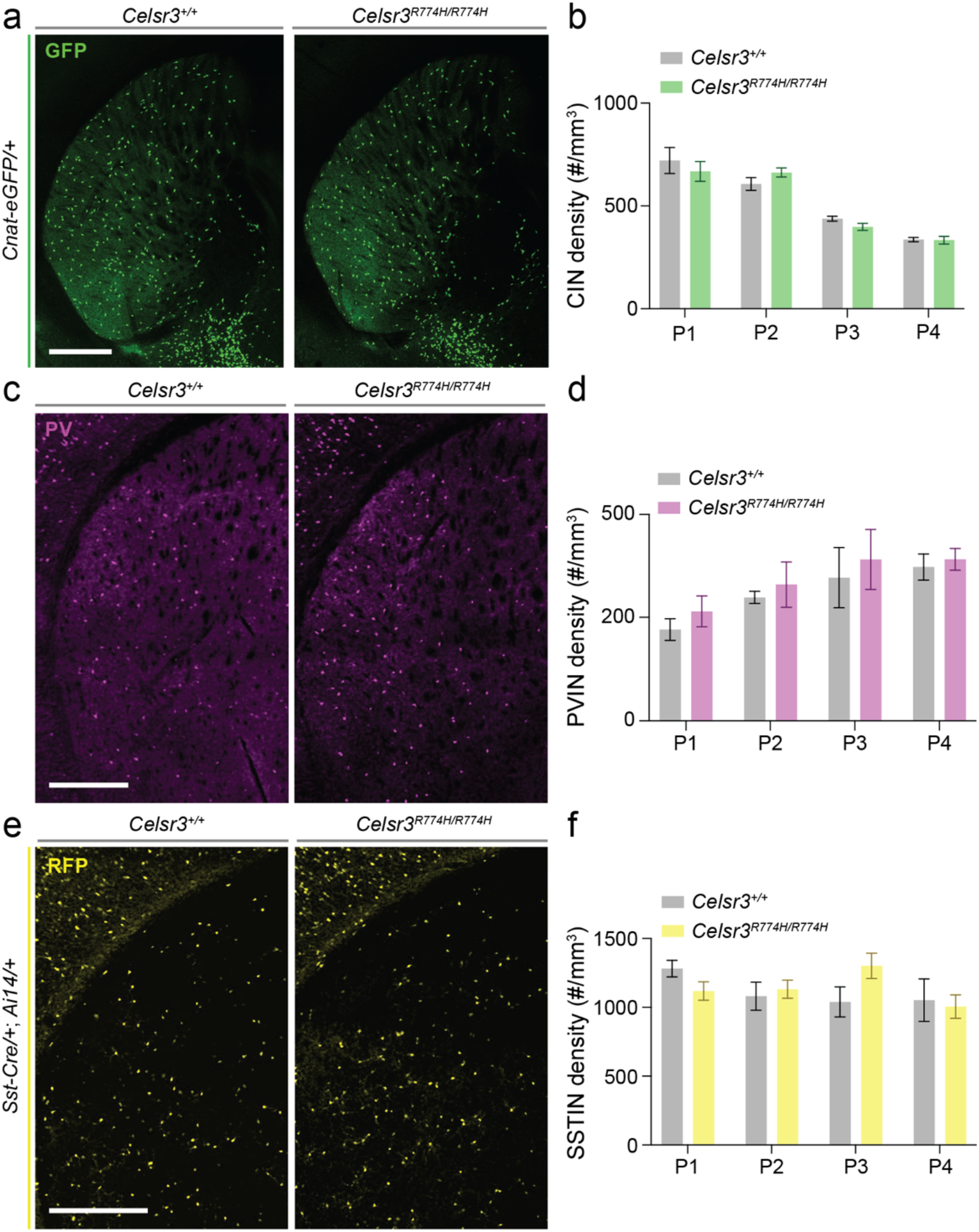
*Celsr3^R774H^*-mutant mice have no detectable loss of cholinergic, parvalbumin-expressing, or somatostatin-expressing striatal interneurons. Representative images of *Celsr3^+/+^; Chat-eGFP* and *Celsr3^R774H/R774H^; Chat-eGFP* striatum (a). Density of GFP+ cholinergic interneurons (CINs) in *Celsr3*^+/+^ (n = 7) and *Celsr3^R774H/R774H^* (n = 10) striatum at four axial positions (p = 0.6728, 2way ANOVA, b). Representative images of PV labelling in the striatum of *Celsr3*^+/+^ and *Celsr3^R774H/R774H^* mice (c). Density of parvalbumin interneurons (PVINs) in the striatum at four axial positions (p = 0.3003, 2way ANOVA, d). Representative images of *Sst-Cre/+; Ai14/+; Celsr3^+/+^* and *Sst-Cre/+; Ai14/+; Celsr3^R774H/R774H^* striatum (e). Density of tdTomato+ somatostatin interneurons (SSTINs_ in the striatum of *Celsr3*^+/+^ (n = 6) and *Celsr3^R774H/R774H^* (n = 5) mice at four axial positions (p = 0.7279, 2way ANOVA, f). Scale bars represent 500 μm.

Our findings in mice suggest human mutations in *CELSR3* may affect the ability of neurons to pattern their dendritic arborizations and receptive fields within CSTC loops (Fig. 3). Deep layer cortical pyramidal neurons in Celsr3^R774H^ animals have atrophic basal dendrites, whereas striatal cholinergic interneurons tend to have more compact arborizations, with more crossings proximal to the soma (Fig. 5). The observation of dendritic patterning changes in cortical pyramidal neurons was an unexpected additional measure that hinged upon germline Cre, which has been previously reported (Luo et al., 2020). Reduced complexity of basal dendrites, however, is in line with previous findings from mice with conditional loss of *Celsr3* in *Dlx5/6-Cre:Celsr3^FLX/FLX^:Thy1-YFP* mice that showed “blunted” dendrites, and also dendritic spine loss in deep layer cortical pyramidal neurons using *Foxg1-Cre* (Zhou et al., 2010). Furthermore, hippocampal CA1 neurons also showed atrophic basal dendrites and loss of dendritic spines in *Celsr3^FLX/FLX^:Foxg1-Cre* mice (Feng et al., 2012). Our results suggest the Celsr3^R774H^ amino acid substitution exerts partial loss-of-function effects on the protein. While we did not detect a significant reduction in overall spine density in cortical pyramidal neurons from Celsr3^R774H^ animals (Fig. 3), we did detect a reduction in spine-like processes on the secondary dendrites of striatal cholinergic interneurons (Fig. 5). We also found changes to the types and distributions of dendritic spines along the secondary basal dendrite of cortical pyramidal neurons (Fig. 3), with a significant loss of stubby spines and a trend towards an increase in long-thin spines. This suggests Celsr3 cadherin repeats are important for regulating the types and distributions of dendritic spines, and that spine maturity or spine turnover may be affected in TD. Examination of long-term potentiation capacity within both cortical pyramidal neurons and striatal cholinergic interneurons may reveal functional impacts that align with these anatomical observations.

**Figure 5.**
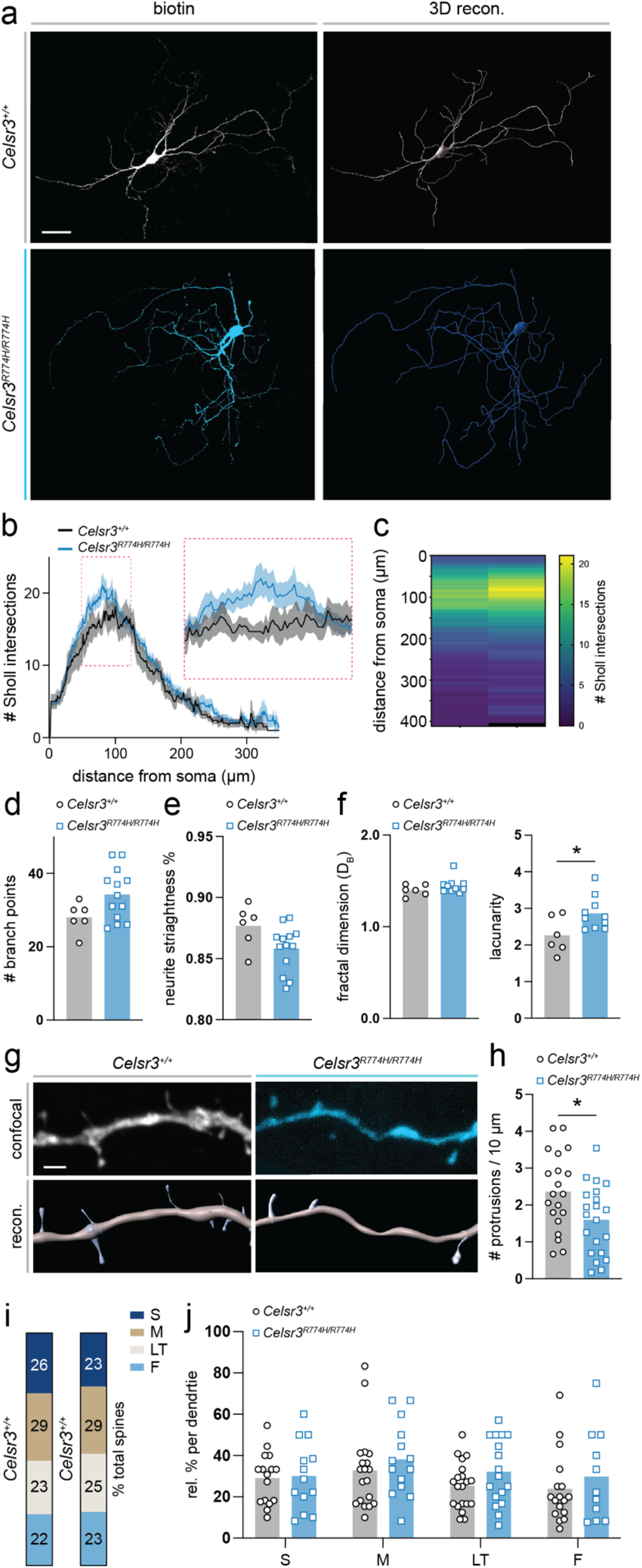
Increased neurite complexity and altered spine-like protrusions in striatal cholinergic interneurons of *Celsr3^R774H^*-mutant mice. Representative images of confocal maximum intensity projections of biotin-filled neurons (left) and their 3D reconstructions (right) in *Celsr3*^+/+^ (top) and *Celsr3^R774H/R774H^* (bottom) mice (a). Scale bar represents 50 μm. Sholl analysis of *Celsr3*^+/+^ (n = 6) and *Celsr3^R774H/R774H^* (n = 13) reconstructed neurons (genotype effect: p < 0.001, 2way ANOVA, b). Inset shows enlargement of Sholl plot ROI (magenta dotted box). Heat map of Sholl intersections vs. distance from soma in *Celsr3*^+/+^ (left) and *Celsr3^R774H/R774H^* mice (right, c). Total number of branch points (p = 0.0619, t-test, d). Neurite straightness score (p = 0.0546, t-test, e). Fractal dimension (p =0.2635, Mann-Whitney test, left) and lacunarity (p = 0.0379, t-test, right) measures (f). 3 neurons were excluded from the *Celsr3^R774H/R774H^* group for this particular analysis due to background pixels that interfered with Db and lacunarity scoring. Representative images of confocal maximum intensity projections of second order dendrite ROIs in *Celsr3*^+/+^ (left, top) and *Celsr3^R774H/R774H^* (right, top) mice and their 3D reconstructions and semiautomatic spine detection (lower panels, g). Scale bar represents 2 μm. Average spine density on second order dendrites (p = 0.0184, t-test, h). Relative contributions (%) of different spine classes on second order dendrites (S = stubby, M = mushroom, LT = long thin, F = filopodia). Relative contributions of spine classes shown per replicate (p = 0.0946, 2way ANOVA, j). Zero values were excluded.

A recent study in rodents suggests cholinergic interneurons show a transient increase in dendritic complexity during the second postnatal week (McGuirt et al., 2021). This period is marked by overgrowth and increased number of crossings, followed by regression starting around the third postnatal week. These findings are interesting because they are similar to those found in Celsr3^R774H^ animals (Fig. 5), suggesting Celsr3 has important functions for dendritic patterning at critical timepoints while cholinergic interneurons are differentiating. Increased lacunarity in *Celsr3^R774H/R774H^* cholinergic interneurons indicates higher levels of heterogeneity in the space-filling properties of their arbors. This could impact how cholinergic interneurons integrate within local striatal circuitry and also shift their electrotonic properties. In addition, we also find less filopodia spine-like appendages on the secondary dendrites of cholinergic interneurons, which normally possess only few spines (Fig. 5) (Poppi et al., 2021). Notably, studies have shown that Celsr3 and PCP signalling is required for excitatory synapse formation in the hippocampus and cerebellum (Feng et al., 2012; Zhou et al., 2021). Thus, perhaps similar to mutations in cell adhesion proteins associated with related neurodevelopmental disorders such as autism spectrum disorder, human mutations in CELSR3 may affect the ability of neurons within CSTC loops to pattern their receptive fields and to form and/or maintain functional synapses.

TD is thought to involve dysregulated striatal dopamine signaling (Rapanelli et al., 2014; Singer et al., 2002; Wong et al., 2008), and drugs that act on D2 dopamine receptor signaling are still a mainstay treatment. Striatal dopamine and acetylcholine work in balance, where high dopamine and low acetylcholine striatal levels are generally associated with hyperkinesia, and conversely, low dopamine and high acetylcholine striatal levels are associated with gait freezing (Barbeau, 1962). Based on this “see-saw” theory and the idea that TD may align with a hyperdopaminergic striatal state, we hypothesized that cholinergic interneurons in *Celsr3^R774H/R774H^* mice would exhibit reduced intrinsic excitability. Surprisingly, we found the opposite. Though mild, *Celsr3^R774H/R774H^* cholinergic interneurons fire APs at a higher frequency than their littermate *Celrs3*^+/+^ controls (Fig. 6), which could be due to either a subtle shift in intrinsic conductances that govern RMP, AP threshold, and AP discharge frequency, or a change in neurite patterning that alters their electrotonic properties (Mainen and Sejnowski, 1996). A more modern take on DA/ACh balance in the striatum is that it is a far more complex interplay with temporal and spatial dimensions (Surmeier and Graybiel, 2012). For instance, coordinated activity of striatal cholinergic interneurons elicits local DA release via excitation of nicotinic receptors on nigrostriatal axons (Cachope et al., 2012; Threlfell et al., 2012). Elevated AP frequency in *Celsr3^R774H/R774H^* cholinergic interneurons could reflect more nuanced changes in D2 and/or muscarinic M2 acetylcholine receptor intracellular signalling arising from changes to the local chemical milieu, and even a small change in cholinergic interneuron firing properties could have effects on tonic and phasic striatal dopamine signalling. These mechanisms are currently being investigated further in the *Celsr3^R774H^* model, as well as in mice carrying other mutations in *Celsr3*.

**Figure 6.**
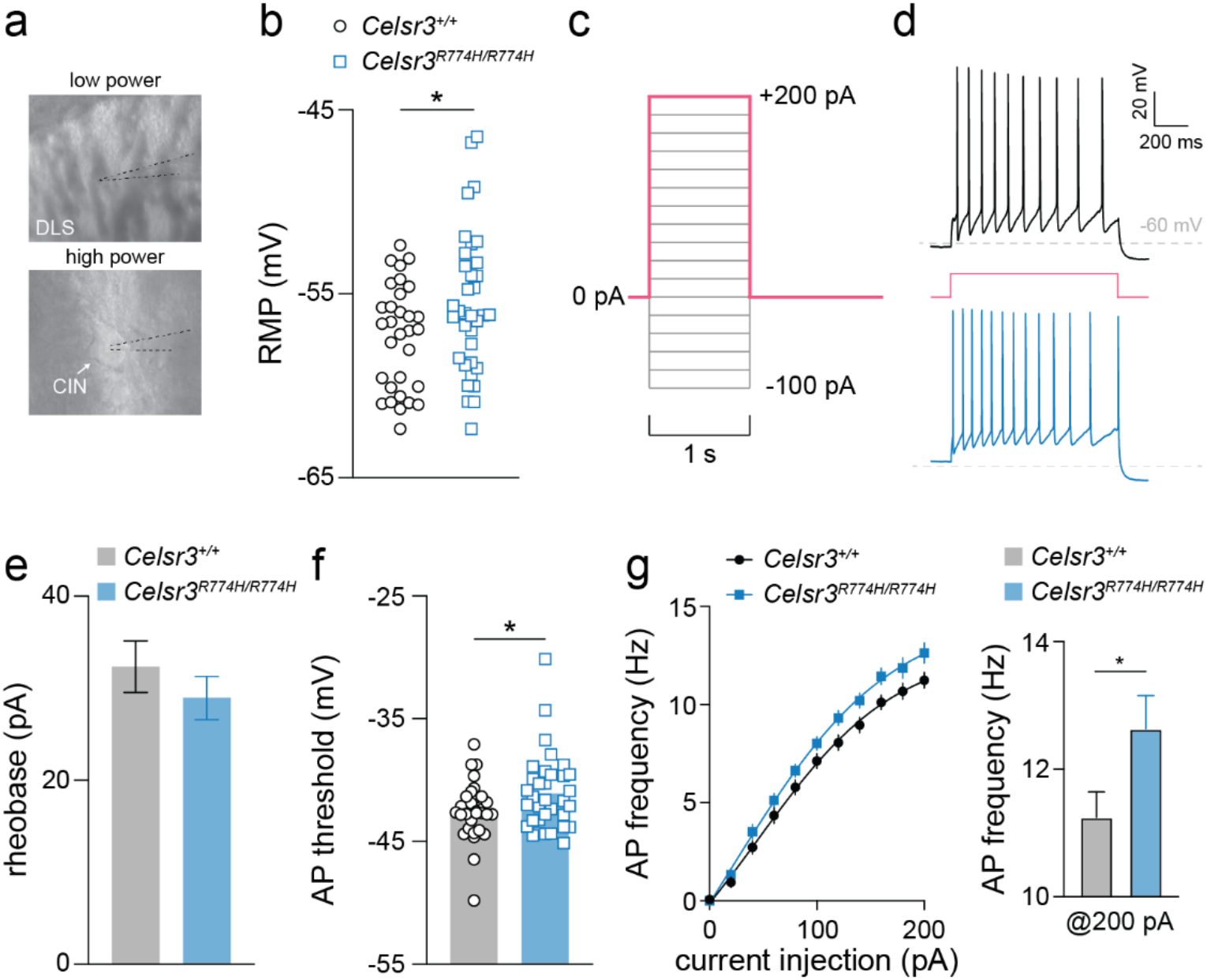
Striatal cholinergic interneurons of *Celsr3^R774H^*-mutant mice show mild intrinsic hyperexcitability. DIC images during recording at low power (top) show placement of electrode in the dorsolateral striatum (DLS) and high power (bottom) showing placement of electrode on an identified cholinergic interneuron (CIN) (a). Resting membrane potential (RMP) of recorded *Celsr3*^+/+^ (n = 31) and *Celsr3^R774H/R774H^* (n = 35) Cholinergic interneurons (p = 0.0386, two-tailed t-test, b). Depolarizing current injection ladder used to characterize evoked action potentials in current clamp mode (c). Representative traces of a *Celsr3*^+/+^ (top, black trace) and a *Celsr3^R774H/R774H^* (lower, blue trace) tonically firing CIN in response to 200 pA current injection (red step, d). Rheobase of *Celsr3*^+/+^ and *Celsr3^R774H/R774H^* Cholinergic interneurons (p = 0.3505, Mann-Whitney test, e). Action potential (AP) threshold of *Celsr3^+/+^* and *Celsr3^R774H/R774H^* Cholinergic interneurons (p = 0.0456, unpaired t-test, f). f/I plot of *Celsr3*^+/+^ (n = 26) and *Celsr3^R774H/R774H^* Cholinergic interneurons (n = 19, left plot, p<0.0001, nonlinear fit - different curve for each dataset). AP frequency @ 200 pA injection for *Celsr3*^+/+^ and *Celsr3^R774H/R774H^* Cholinergic interneurons (right graph, p = 0.0382, two-tailed t test).

It is somewhat unexpected that Celsr3^R774H^ animals exhibit only very mild changes to motor behavior, and do not show obvious tic-like stereotypies at baseline. In agreement, however, *Hdc* knock-out animals also do not show tic-like stereotypies at baseline without stress or amphetamine challenge (Castellan Baldan et al., 2014), consistent with findings that stress and sensory overload can exacerbate tics in humans. Moreover, targeted ablation of cholinergic or parvalbumin interneurons in the dorsal striatum, mimicking striatal interneuron loss found in some humans, only leads to tic-like stereotypies following acute stress or amphetamine challenge (Rapanelli et al., 2017b; Xu et al., 2015; Xu et al., 2016). Ablating ~50% of both populations simultaneously causes spontaneous stereotypies in males but not females (Rapanelli et al., 2017b), whereas more extensive loss of cholinergic interneurons can cause compulsive-like social behavior and repetitive digging (Martos et al., 2017). We also find that Celsr3^R774H^ homozygous females, but not males, bury more marbles (Fig. 7), consistent with changes to compulsive or perseverative behaviors in response to environmentally driven stimuli. Interestingly, studies suggest that complex tics, which can reflect compulsions and are often performed in a ritualistic manner, are more common in females with TD versus males (Garris and Quigg, 2021; Hirschtritt et al., 2015). Thus, our genetic models may be useful for modelling sex-specific behavioral differences in TD. Finally, using motion sequencing to parse motor behavior into discrete modules, and computational approaches that use natural language processing to examine how short motion sequences are embedded into groupings, we detect subtle but distinct changes to action selection in Celsr3^R774H^ animals (Fig. 7). This suggests the Celsr3^R774H^ amino acid substitution exerts mild effects on motor functions that may become more apparent with stress, anxiety, or sensory overload. It will also be interesting to apply machine learning approaches that can analyze facial movements (Dolensek et al., 2020), as facial tics are more common in TD and may have been missed in our models.

**Figure 7.**
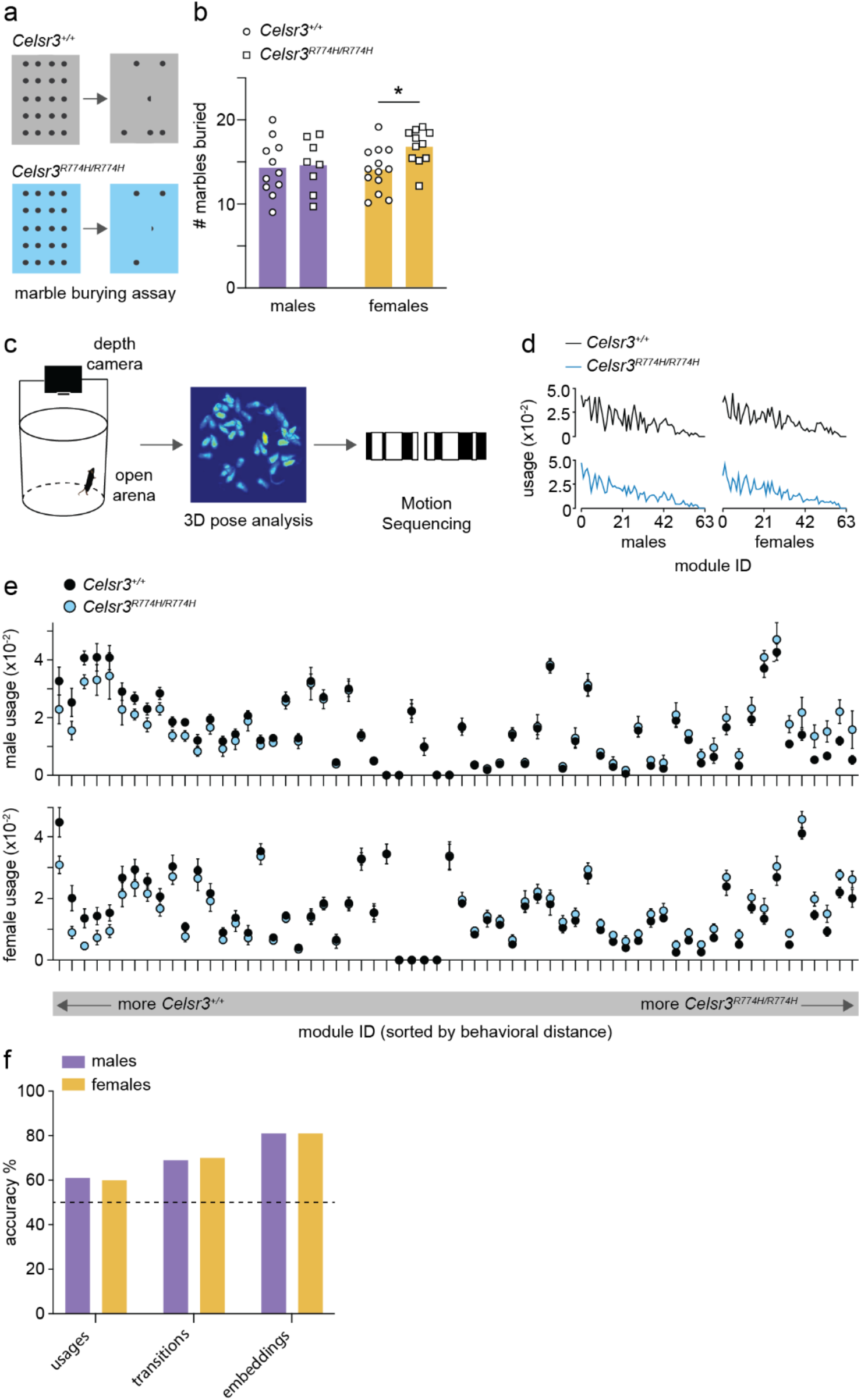
*Celsr3^R774H^*-mutant mice show perseverative tendencies and natural language processing can predict genotype to 81% accuracy. The marble burying assay consists of 20 glass marbles arranged in a 4 x 5 grid on top of 5 cm chip bedding (a). The average number of marbles buried across three trials by *Celsr3*^+/+^ and *Celsr3^R774H/R774H^* mice in 30 mins (b). Female *Celsr3^R774H/R774H^* mice (n = 11) buried 16.66 +/− 0.64 marbles compared to *Celsr3^+/+^* littermates (n = 13) who buried 13.92 +/− 0.72 marbles (p = 0.011, unpaired t-test; b). Mouse behavior was recorded in an open arena with a depth camera, and behavioral features were sequenced using unsupervised machine learning (Motion Sequencing, c). Usage of behavioural modules (ordered 0-63) is similar when comparing across genotype in males (left graphs) and females (right graphs, d). Module usage with modules ordered by behavioral distance (left x-axis – *Celsr3*^+/+^ usage tends to be higher, right x-axis – *Celsr3^R774H/R774H^* usage tends to be higher, e). A comparison of accuracy levels in predicting genotype (*Celsr3*^+/+^, *Celsr3*^*R774H*/+^, or *Celsr3^R774H/R774H^)* based on module usages, transitions between modules, and learned embeddings (f).

In summary, our findings in Celsr3^R774H^ animals point to subtle but detectable changes to the ability of neurons to pattern their receptive fields within CSTC loops, and also the ability of neurons in both the cortex and striatum to regulate the types and distributions of dendritic spines. This suggests that human mutations in *CELSR3* cause TD by affecting how neurons integrate and signal within CSTC circuits, rather than causing cell loss or other types of structural brain abnormalities. It will be important to confirm and extend our findings in other genetic models engineered to express mutations in different functional domains of Celsr3. It will also be interesting to compare neuronal and behavioral phenotypes with models engineered to express human mutations in *WWC1* and *OPA1*, work that is currently ongoing.

---

**Supplementary Table S1.** Mouse lines.

| Line name | Description/Use | Supplier | Stock# |
| --- | --- | --- | --- |
| *A2a-Cre* | Cre recombinase expressed under control of *A2a*, used to visualize indirect pathway axons in mouse brain | MMRRC | 036158-UCD |
| *Ai14* | Reporter line that expresses TdTomato in Cre recombinase expressing cells, used to visualize direct and indirect pathway axons when crossed with *Drd1-Cre* and *A2a-Cre*, respectively, and used to quantify density and nearest neighbor distribution of cortical and striatal SSTINs | JAX Mice | 007914 |
| *C57BL/6* | Wild type inbred line used as a background strain, and for backcrossing + line refreshing | JAX Mice | 000664 |
| *Celsr3-eGFP* | Knock-in eGFP line used to study *Celsr3* expression patterns in mouse brain | Mario Cappechi, University of Utah (Ying et al., 2009) | RRID:MGI:3849330 |
| *Celsr3<sup>R774H</sup>* | Line carrying point mutation in *Celsr3*, used in all experiments | generated in house | n/a |
| *Chat-eGFP* | BAC transgenic line expressing eGFP in cholinergic cells, used to quantify density of striatal Cholinergic interneurons + for targeted recordings in dorsolateral striatum | JAX Mice | 007902 |
| *Drd1-Cre* | Cre recombinase expressed under control of *Drd1*, used to visualize direct pathway axons in mouse brain | MMRRC | 030989-UCD |
| *Pvalb-Cre* | Cre recombinase expressed under control of *Pvalb*, used for off-target sparse cell labelling of layer V cortical neurons | JAX Mice | 012358 |
| *Sst-Cre* | Cre recombinase expressed under control of *Sst*, used to quantify cortical + striatal SSTIN density | JAX Mice | 028864 |

**Supplementary Table S2.** Criteria for spine classification.

| Spine Class | Criteria |
| --- | --- |
| **Stubby** | Spine length < 1 μm |
| **Mushroom** | Spine length < 3 μm and spine head width > spine neck width*2 |
| **Long Thin** | Spine head width ≥ spine neck width |
| **Filopodia** | True |

## Supplementary Methods

### Immunofluorescent labelling

The typical protocol for immunofluorescent labelling consisted of 0.1 M PBS washes, followed by a 1-hour incubation in either normal donkey serum or normal goat serum, depending on the host species of secondary antibodies used. This was followed by incubation in primary antibody solution, washes in 0.1 M PBS, incubation in secondary antibody solution, washes in 0.1 M PBS, and finally mounting onto glass microscope slides (VWR) using Fluoromount-G mounting media (Southern Biotech). All anatomy data were acquired using confocal microscopy (Zeiss LSM 700 or Zeiss LSM 800) except for direct and indirect pathway visualization studies where data were collected on a Leica M165FC stereomicroscope with CoolLED illumination.

### Accelerated rotarod test

The Rota-rod apparatus (LE8205, Panlab) was used to assess motor learning capabilities of mice. Mice were placed on the rod before the test started and the rod accelerated from 4 rpm to 40 rpm in 5 minutes. The duration of time that mice could stay on the rotating rod in each trial (latency) was recorded automatically. Mice were given at least 1 min recovery time between trials. During testing, the rod was kept dry and clean. Mice were tested for 5 trials per day over 6 consecutive days. The average of latency each mouse per day was plotted.

### SmartCage testing system

SmartCage (AfaSci) equipped with base and upper layers of infrared (IR) beam arrays was used to analyze mouse locomotion. Mice were habituated for at least one hour with dim light. Mice were then placed in the homecage sized SmartCages and were left to freely behave for 30 minutes. Mouse movement blocked the IR beams and signals were recorded automatically. Data were binned into 10-minute time blocks.

**Extended Data 1.**
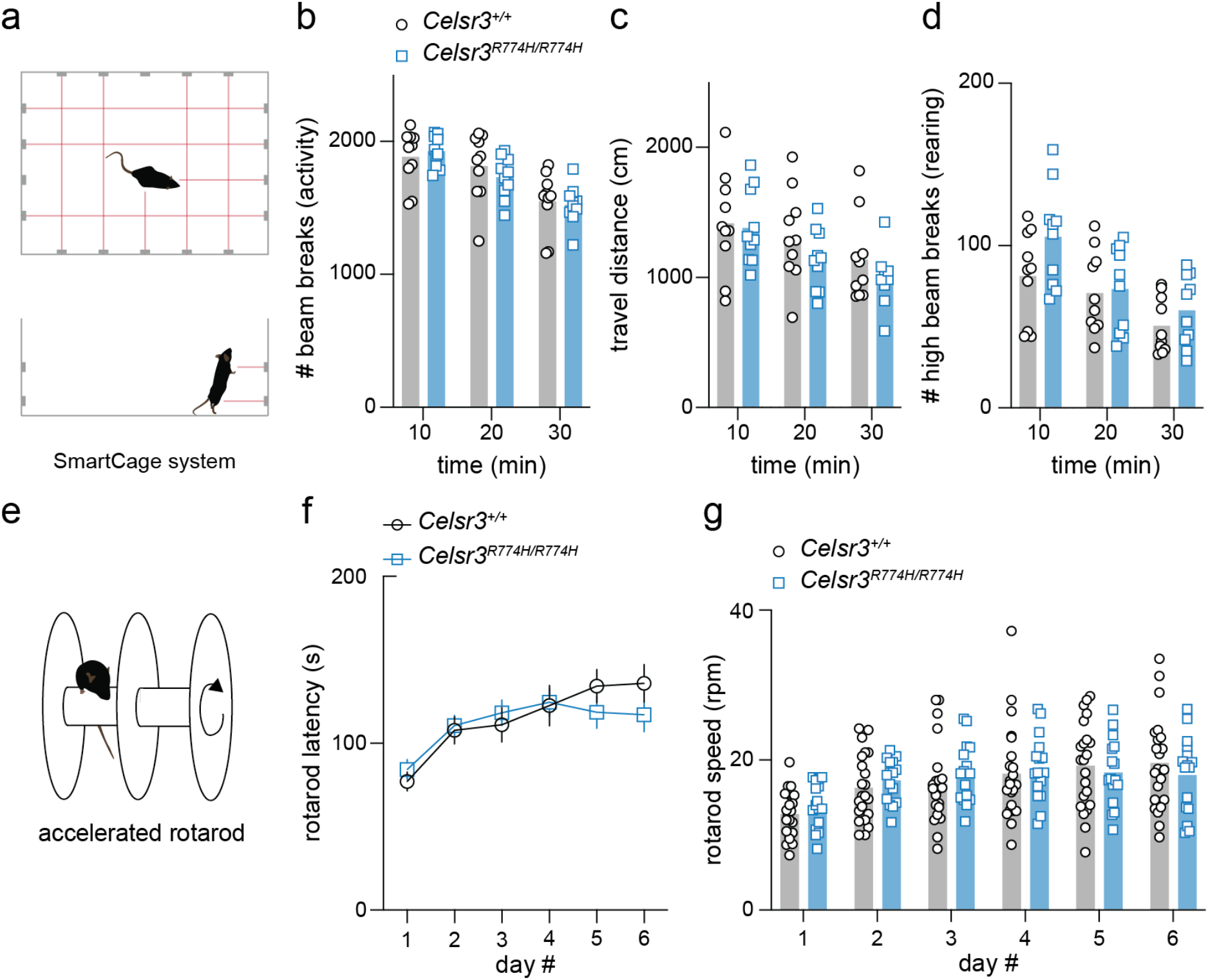
*Celsr3^R774H^*-mutant mice do not exhibit hyperactivity or motor learning deficits. Mice were placed in a SmartCage system fitted with a lower and upper IR beam array to measure activity and rearing behavior (a). Total number of beam breaks within 10-minute windows for *Celsr3*^+/+^ (n = 10) and *Celsr3^R774H/R774H^* mice (n = 10, p = 0.6249, 2way ANOVA, b). Total travel distance for *Celsr3*^+/+^ and *Celsr3^R774H/R774H^* mice over 30 minutes (p = 0.1323, 2way ANOVA, c). Upper beam breaks (rearing activity) of *Celsr3*^+/+^ and *Celsr3^R774H/R774H^* mice (p = 0.0717, 2way ANOVA). Mice were assessed on an accelerated rotarod test (e). Average latency to fall from rotarod vs. day of testing for *Celsr3*^+/+^ (n = 20) and *Celsr3^R774H/R774H^* mice (n = 18, p = 0.6300, 2way ANOVA) (f). Maximum rotarod speed (revolutions per minute) vs day of test (p = 0.6760, 2way ANOVA, g).

**Extended Data 2.**
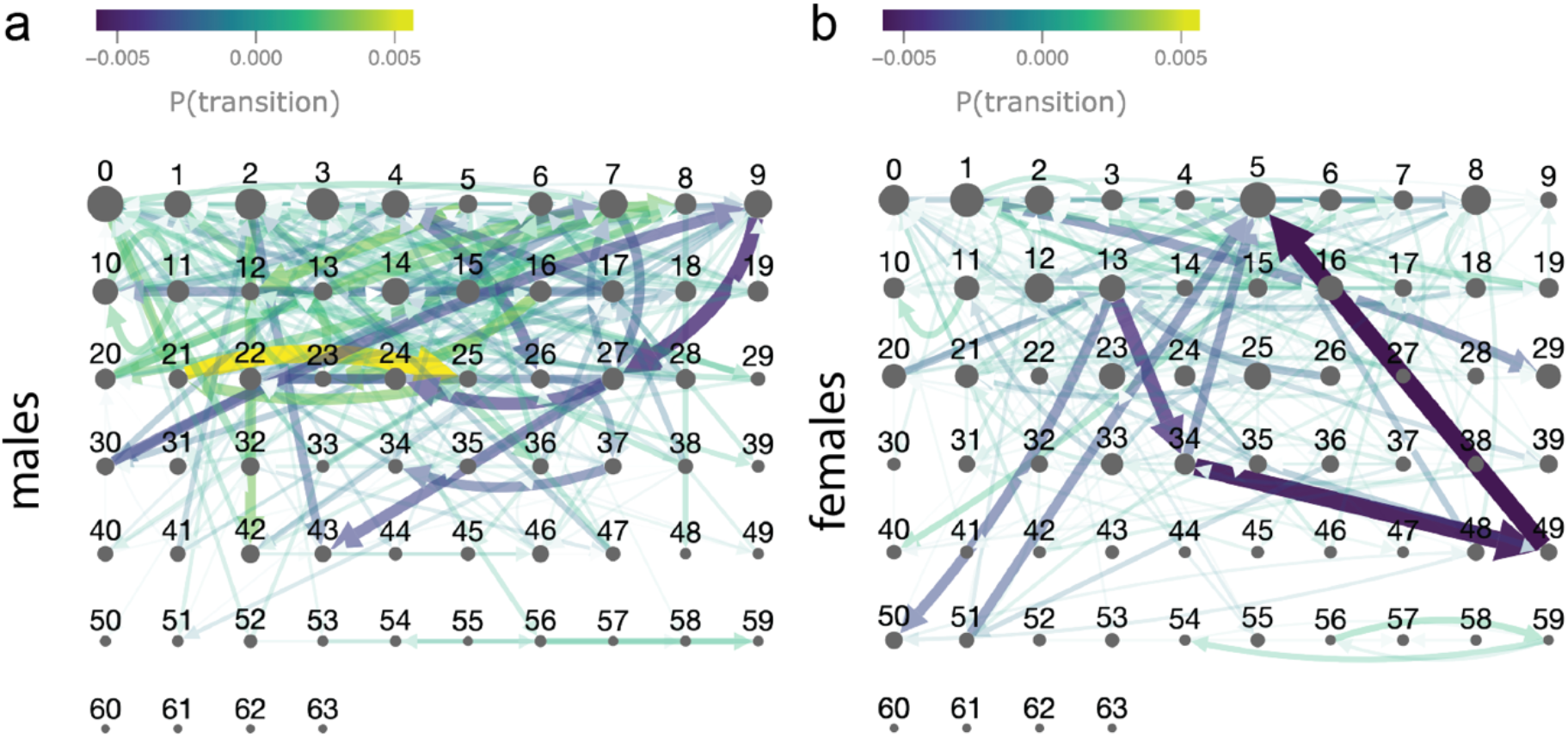
Altered probabilities of transitioning between behavioral modules in *Celsr3*^+/+^ and *Celsr3^R774H/R774H^* mice. Transition probability matrix for *Celsr3^R774H/R774H^* males (n = 8) relative to *Celsr3*^+/+^ males (n = 12, a). Transition probability matrix for *Celsr3^R774H/R774H^* females (n = 13) relative to *Celsr3*^+/+^ females (n = 14). Pruning threshold was set to p = 0.0001. Dot size is proportional to module usage.

**Extended Data 3.**
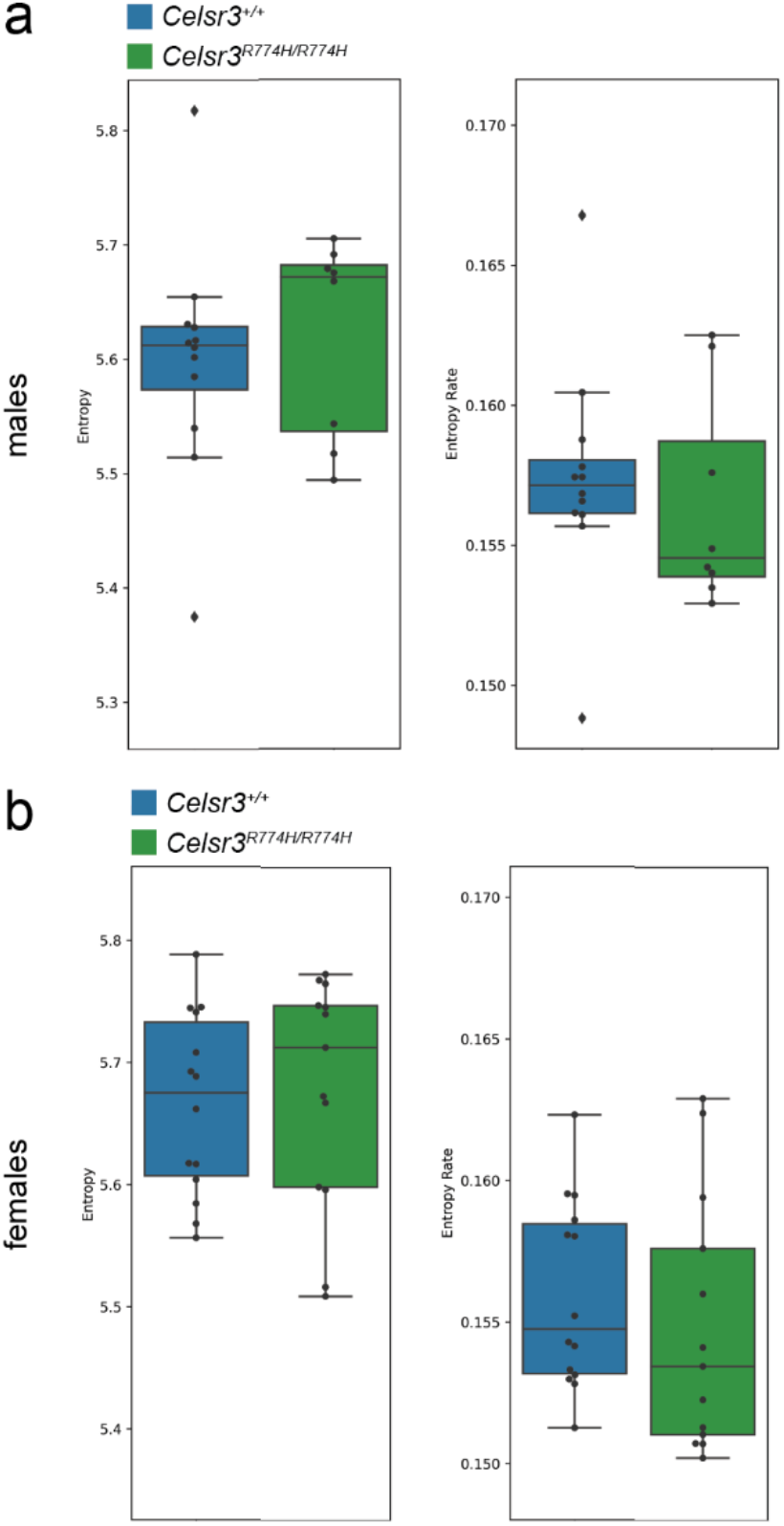
Entropy plots for *Celsr3*^+/+^ and *Celsr3^R774H/R774H^* mice. Entropy plot (left) and entropy rate plot (right) for *Celsr3*^+/+^ and *Celsr3^R774H/R774H^* males (a). Entropy plot (left) and entropy rate plot (right) for *Celsr3*^+/+^ and *Celsr3^R774H/R774H^* females (b).

**Extended Data 4.**
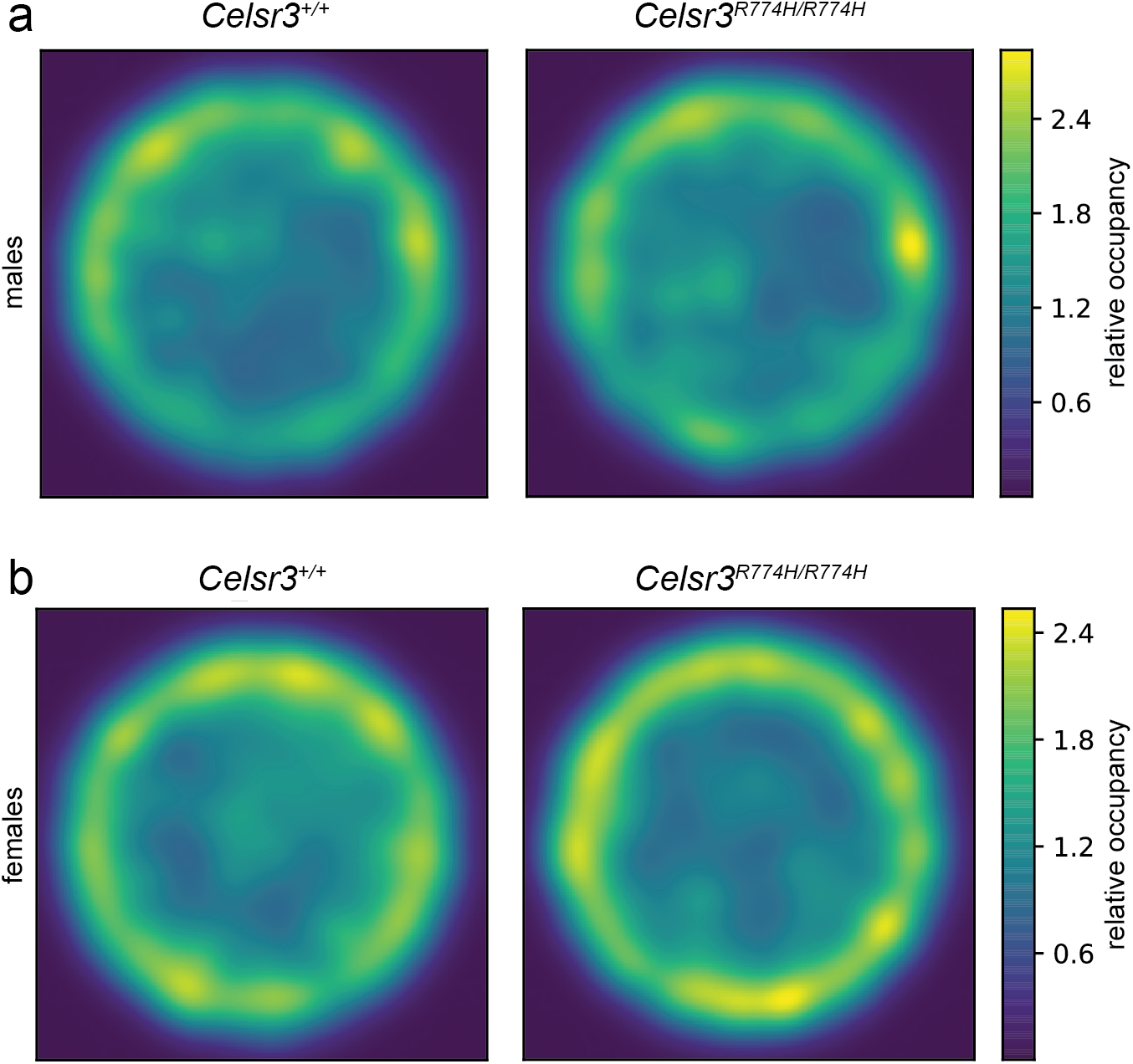
Occupancy heat maps for *Celsr3*^+/+^ and *Celsr3^R774H/R774H^* mice in open arena. Relative occupancy heatmaps for *Celsr3*^+/+^ and *Celsr3^R774H/R774H^* male mice (a). Relative occupancy heatmaps for *Celsr3*^+/+^ and *Celsr3^R774H/R774H^* female mice (b).

## Acknowledgements

We would like to thank A.K. for their valuable contributions to the early stages of this study.

